# Hypersensitive Detection of Neurotransmitters in Biological Media by Optically Enhanced Benchtop NMR Spectroscopy

**DOI:** 10.64898/2026.09.11.751013

**Authors:** Anubhab Halder, Samuel C. Carey, Ji Ho Jeong, Ummay Mahfuza Shapla, Catherine F.M. Clewett, Silvia Cavagnero

**Author notes:** Equal contributors (Anubhab Halder, Samuel C. Carey).

## Abstract

Liquid-state Nuclear Magnetic Resonance (NMR) is an invaluable tool to gain atomic-resolution insights onto molecular structure and dynamics. The impact of NMR spectroscopy, however, is curtailed by high costs and hard-to-maintain high-field magnets. While low-field benchtop NMR spectrometers may address the challenge, severe trade-offs in sensitivity and resolution limit the appeal of this avenue. Here, we introduce a set of novel optically enhanced NMR pulse sequences and procedures (hyperpolarization toolkit) to readily generate ^13^C and ^1^H nuclear-spin hyperpolarization *in situ* on benchtop spectrometers. This approach leads to unprecedented sensitivity gains, enabling atomic-resolution conformation-dependent detection of a broad range of aromatic compounds, including clinically relevant metabolites and biomarkers. Neurotransmitters including epinephrine, serotonin and melatonin are detected at nanomolar/micromolar levels by 1D/2D photochemically-enhanced benchtop NMR in buffer and physiological media including human serum. This advance propels benchtop NMR from a predominantly pedagogical tool to a powerful bioanalytical resource for efficient neurotransmitter identification.

## Introduction

Many clinically important biomolecules (e.g., amino acids, metabolites, pharmaceuticals) are present at low micromolar to picomolar concentration in the human body. Among them, amino acids and their catabolic products play key roles in signaling, metabolism, and disease progression.^1–3^ The accurate detection of metabolites in unmodified biofluids (e.g., serum, urine) is vital for biomedical research, diagnostics, and pharmaceutical sciences.^4,5^ While detection methods including liquid-chromatography coupled to mass spectrometry (LC-MS),^6^ fluorescence and biochemical assays^7–9^ can reach nM-pM sensitivity, these approaches typically lack information on the conformational properties of target molecules, and often require complex sample-preparation steps involving ionization or purification, thus preventing *in situ* analysis.^10^ Note that hydrogen-deuterium exchange mass spectrometry (HDX-MS), which provides indirect shape information on macromolecules, does not work for small molecules, given their negligible extent of protection from H/D exchange upon transfer in deuterated media. Hence, it is desirable to develop alternative technologies that enable high-resolution, high-sensitivity, inexpensive and shape-dependent *in situ* analysis of small biomolecules in complex physiological media.

Solution-state NMR spectroscopy fulfils most of the above requirements. However, it is an inherently low-sensitivity technique. Thus, there is a critical need for advances that enhance the detection thresholds of NMR spectroscopy and enable bioanalytical investigations at low sample concentration (≤ micromolar). Upon driving nuclear spins into a transient non-equilibrium condition that perturbs equilibrium populations **(Fig. 1A)**, it is possible to develop hyperpolarized states that lead to improvements in NMR sensitivity by orders of magnitude.^11,12^ Several hyperpolarization approaches including photochemically induced dynamic nuclear polarization (photo-CIDNP), parahydrogen induced polarization (PHIP),^13,14^ optical pumping (OP)^15^ and dissolution dynamic nuclear polarization (dDNP),^16,17^ have been devised to mitigate the low sensitivity challenges of NMR in liquids. The photo-CIDNP optically enhanced hyperpolarization technique is characterized by high simplicity and low cost, and it is compatible with mild physiologically relevant conditions. In addition, photo-CIDNP generates rapid *in situ* hyperpolarization (∼0.2 seconds) and it has multiple-transient data-collection capabilities, which in turn enable both 1D and multidimensional (2D, etc.) experiments.^18,19^ Further, recent methodological developments extended the applicability of photo-CIDNP to the low-concentration regime via an approach known as low-concentration photochemically induced dynamic nuclear polarization (LC-photo-CIDNP).^20–22^ This approach employs oxygen-scavenging enzymes and dyes with long photoexcited-state lifetimes (**Fig. 1B**), to facilitate productive photoexcited-dye/molecule-of-interest collisions even at low sample concentration.

**Figure 1.**
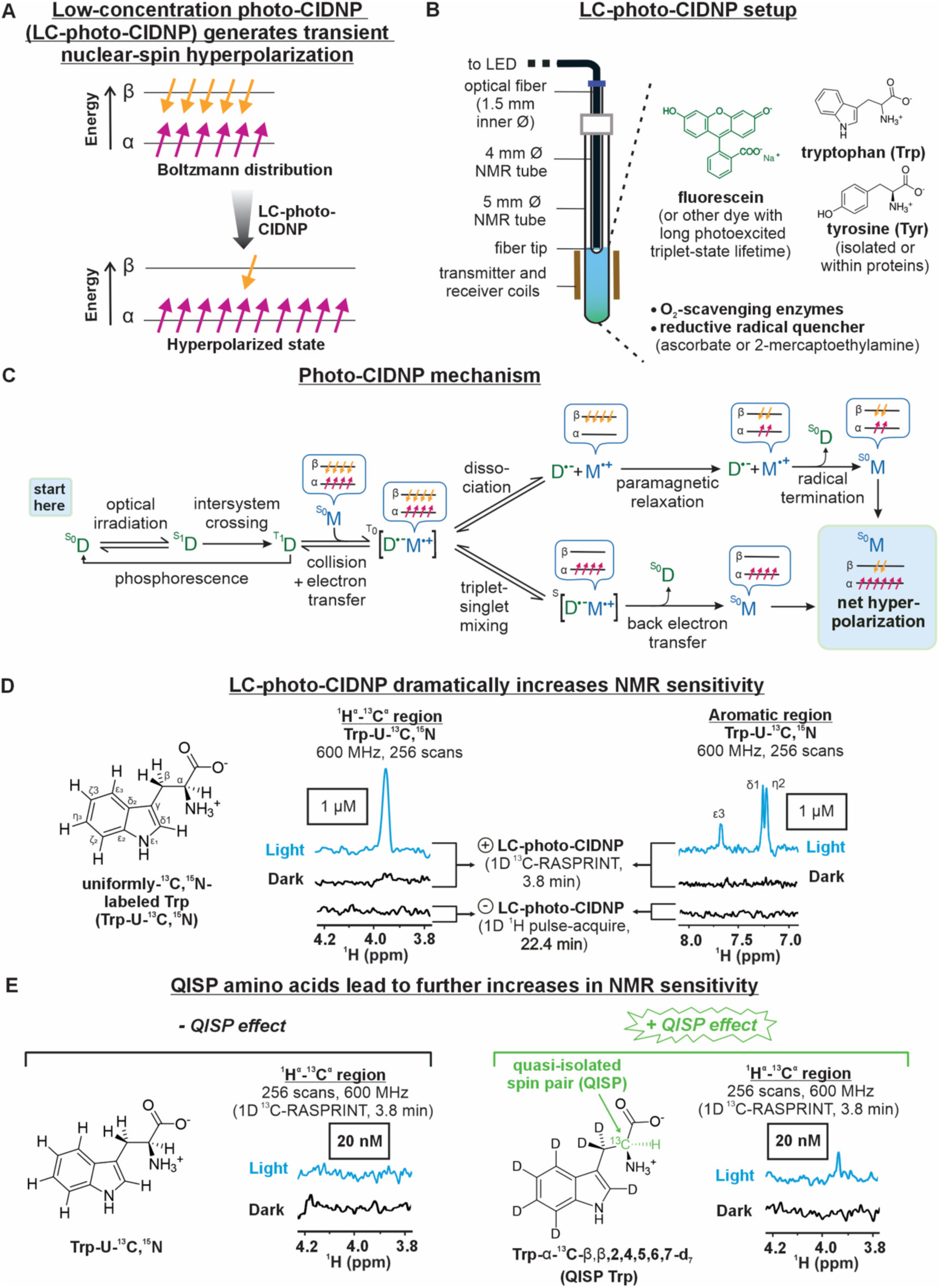
Low-Concentration photochemically induced dynamic nuclear polarization (LC-photo-CIDNP) generates unprecedented NMR sensitivity. **A.** Schematic representation of nuclear spin hyperpolarization in LC-photo-CIDNP. **B.** Cartoon illustrating a LED-enhanced LC-photo-CIDNP NMR apparatus (see Methods).^12^ **C.** Schematic representation of major mechanistic steps underlying liquid state photo-CIDNP (or LC-photo-CIDNP). **D.** 1D ^13^C RASPRINT spectra under light (LED-on) and dark (LED-off) conditions and comparison to a conventional pulse-acquire (1D ^1^H NMR) of a 1 μM Trp-U-^13^C, ^15^N in aqueous buffer. The pulse-acquire experiments were performed with W5 WATERGATE water suppression pulse sequence with excitation sculpting and GARP ^13^C decoupling during acquisition (n = 2). **E.** 1D ^13^C RASPRINT spectra under light (LED-on) and dark (LED-off) conditions of a 20 nM QISP Trp in aqueous buffer. All ^13^C RASPRINT LC-photo-CIDNP spectra shown here were in 10 mM phosphate buffer (pH ∼7.2), used 2.5 μM fluorescein as the photosensitizer dye and were collected at 24°C with a 0.05 s recycle delay and 0.2 s LED irradiation time per scan (n = 2). 1D ^1^H NMR spectrum was collected with a 5 s recycle delay. All data were collected on a 14.1 T (600 MHz) NMR spectrometer.

Despite the above advantages, the cost of the purchase and maintenance of high-field NMR spectrometers has recently become prohibitive, due to pricy hardware components and to the decreased worldwide availability of cryogens, especially helium, employed for the routine cooling of high-field superconducting magnets.^20,21^ In all, the routine atomic-resolution analysis of metabolites by hyperpolarization-enhanced NMR spectroscopy would greatly benefit from a lower-cost/lower-maintenance version of this technique.

Cryogen-free benchtop NMR spectrometers, which are typically equipped with permanent magnets and room-temperature (i.e., non-cryogenic) probes, offer an attractive alternative.^25,26^ These compact low-field systems are less sensitive than high-field spectrometers, typically operate between 42 and 125 MHz (0.99-2.94 T), and are much smaller and much less expensive than traditional high-field instruments (e.g., ≥400 MHz, 9.40 T). In recent times, the dissolution dynamic nuclear polarization (dDNP) and signal amplification by reversible exchange (SABRE) approaches were proposed to address the low sensitivity of benchtop NMR spectrometers by coupling low-field magnets with nuclear-spin hyperpolarization (**Supplementary Table 1**). Unfortunately, these implementations either work only on a restricted set of molecules (often in non-aqueous solvents) or require cryogenic cooling and sample transfers between apparatuses.^23–26^ In addition, these approaches are 1-scan techniques and are therefore not amenable to multidimensional data collection. In recent studies, the LC-photo-CIDNP NMR technology has also been employed on benchtop NMR spectrometers (**Supplementary Table 1**). However, its range of applicability was not examined in detail, and its promise was not fully exploited.

LC-photo-CIDNP is particularly convenient because, upon choosing appropriate dyes, it can be applied to a wide variety of aromatic-containing molecules. In addition, this technology involves straightforward sample handling, works well with low-concentration molecules of interest, and is compatible with biological media and with multiple-scan acquisitions. Most importantly, LC-photo-CIDNP is theoretically predicted to achieve higher hyperpolarization at lower magnetic fields in the presence of most known dyes, and it is therefore ideally suited to benchtop NMR spectroscopy.^18^ Further, the isotopic-substitution advantage of LC-photo-CIDNP, as shown in **Fig. 1E** for a tryptophan target molecule bearing a quasi-isolated spin pair (QISP Trp), enables ready detection at 20 nM levels.^27^ Hence, in summary, there is a compelling need to further enhance the capabilities of LC-photo-CIDNP benchtop NMR for bioanalytical purposes.

In this work we develop a toolkit of novel pulse sequences and experimental procedures suitable to perform low-field (80 MHz) LC-photo-CIDNP benchtop NMR on a wide range aromatic-ring containing metabolites and biopharmaceuticals. Clear detectability, down to 100 nM levels, is shown for a variety of clinically relevant neurotransmitters, in both buffered solutions and complex biological media. First, we provide robust theoretical and experimental validation of sensitivity enhancements in two systems: (1) QISP Trp via ¹³C RASPRINT, and (2) unlabeled Trp via ¹H-photo-CIDNP, under physiologically relevant conditions. We achieved an unprecedented detection limit of 2 μM with QISP Trp in ∼ 1 min at 80 MHz (1.88 T). Importantly, we then extend our analysis to a variety of metabolites, neurotransmitters and pharmaceuticals at natural isotopic abundance (e.g., serotonin at 100 nM), collecting atomic-resolution data in ≤ 7 min. This technology is shown to save experimental time by orders of magnitude. QISP Trp shows an LC-photo-CIDNP enhancement factor ε >3000, which, to the best of our knowledge, is the highest **ε** on a benchtop NMR at low-μM concentrations to date. In addition, we demonstrate that LC-photo-CIDNP can be successfully performed in crowded and heterogeneous media including cellular environments and unaltered eukaryotic biofluids (e.g., human serum). In the case of human serum, benchtop LC-photo-CIDNP offers remarkably background-free and unambiguous atomic-resolution detection of neurotransmitters including epinephrine at 10 μM levels, in contrast to high-field. Finally, to improve spectral resolution at 80 MHz, we employ, for the first time, two-dimensional COSY-LC-photo-CIDNP and achieve facile detection of epinephrine at ≥10 μM levels within minutes including polarization transfer to traditionally non-photo-CIDNP-active resonances. In all, our approach enables sophisticated NMR analysis of biomolecules bearing aromatic groups at low field, enabling the facile hypersensitive detection of metabolites, neurotransmitters and pharmaceuticals coupled with atomic-resolution conformational analysis.

## Results

### Research rationale and design principles

Both photo-CIDNP and LC-photo-CIDNP exploit a spin-sorting process (**Fig. 1C**) that proceeds via transiently generated triplet radical pairs. The latter are formed through photoinduced electron transfer between a photosensitizer dye and the target molecule. Spin sorting, when applied to cyclic reactions and in the presence of unpaired-electron-induced paramagnetic relaxation, leads to transient non-Boltzmann populations of nuclear spins, resulting in enhanced NMR sensitivity (**Fig. 1C).**^12,27^ In addition, LC-photo-CIDNP can be coupled with homo- and heteronuclear correlation spectroscopy to enable the direct detection of hyperpolarized ^1^H, ^13^C, ^15^N.^28–30^ For instance, ¹H-detected ¹³C-photo-CIDNP (¹³C RASPRINT) experiments yield ∼1,300-fold sensitivity enhancements, pushing sample detectability far beyond the capabilities of conventional NMR (**Fig. 1D**).^20,30–32^ To further exploit the detection limits of ¹³C RASPRINT, aromatic amino acids carrying quasi-isolated spin pairs (e.g. QISP Trp/Tyr) were designed to reach 20 nM sensitivity. This advance enables ¹H-photo-CIDNP detection of low-μM to low-nM small molecules at natural abundance on high-field and ultra-high-field spectrometers operating at 600 MHz (14.1 T, **Fig. 1E**)^19,33^ and 1.1 GHz (25.9 T),^34^ respectively. Further, LC-photo-CIDNP studies performed in the presence of field cycling showed that QISP amino-acid isotopologs experience higher hyperpolarization enhancements at progressively low applied field, down to 50 MHz (1.17 T).^35^ The latter results showed that low-field hyperpolarization can be advantageous, although some losses due to the mechanical setup were present, and despite the fact that the field-cycling setup is rather complex and not readily available. Given the above findings, we elected to employ existing LC-photo-CIDNP photosensitizers and simple benchtop NMR spectrometers to explore whether ^1^H and ^13^C LC-photo-CIDNP can be developed into a generally applicable technology to detect aromatic-containing bioanalytes in solution at low applied field.

Further, we noted that most neurotransmitters bear one or more aromatic rings, and have redox potentials^36^ and hyperfine coupling constants^37^ extremely compatible with photo-CIDNP. For instance, the standard-state redox potential (E_0_) of serotonin is ca. 0.65 V in water (relative to the standard hydrogen electrode, SHE),^38^ as opposed to tryptophan (Trp), whose corresponding value is E_0_ = 1.015 V (relative to SHE).^21^ Hence, serotonin is estimated to be a more easily oxidized by the photoexcited triplet state of the fluorescein photosensitizer (E_0_ = 1.16 V). Therefore, it is expected to be an even better photo-CIDNP substrate than Trp, in terms of its redox characteristics, if all other factors were equal. In general, several neurotransmitters are also known to endogenously undergo reversible redox reactions in the brain, often involving oxidized radicals.^39^ Hence, they are known to be redox-active compounds, and we reasoned that this class of molecules bears a particularly promising potential to be an excellent target for high-sensitivity and high-resolution *in situ* bioanalytical characterization by LC-photo-CIDNP on benchtop NMR spectrometers.

### Theoretical predictions and experimental validations reveal dramatic ^1^H-detected ^13^C hyperpolarization at low magnetic field

We started with indole-containing model compounds, and employed the theoretical treatment by Adrian^40^ to estimate the applied-field (B_0_) dependence of the expected ^13^C^α^ polarization of two selectively labeled Trp isotopologs, QISP Trp and Trp-U-^13^C, ^15^N (**Fig. 1A**). Note that QISP aromatic-amino-acid isotopologs (QISP Trp/Tyr) retain the large hyperfine coupling constant (HFC) of the ¹³C^α^ nucleus (relative to ^1^H^α^, which has HFC ∼ 0 mT, also see **Supplementary Table 2**) within the photo-CIDNP radical pair. In addition, they minimize the polarization-lowering effects arising from neighboring NMR-active nuclei with large gyromagnetic ratios.^19,41^ The results of the simulations, shown in **Fig. 2C**, highlight that QISP Trp is expected to display a ca. 10-fold higher geminate polarization than Trp-U-¹³C, ¹⁵N, on a benchtop NMR spectrometer operating at 80 MHz. This large increase outcompetes the prediction at 600 MHz, where only ca.1.3-fold greater geminate polarization is expected **(Fig. 2C).** This strong dependence of QISP-like isotopic substitutions at low field arises from the relative magnitude of the electronic Zeeman (ΔgβₑB₀/ħ) and HFC terms in the theoretical expressions for the frequency of triplet-singlet mixing (μ_TS,α,χ_ or μ_TS,β,χ_) for the nuclear-spin state (either α or β) of the nucleus of interest.^42^ In short, at lower B_0_ the Zeeman term becomes smaller and more comparable in magnitude to the HFC term, leading to a larger difference in μ_TS,α,χ_ and μ_TS,β,χ_, leading to higher hyperpolarization. This topic is further discussed in the **Supplementary Information**.

**Figure 2.**
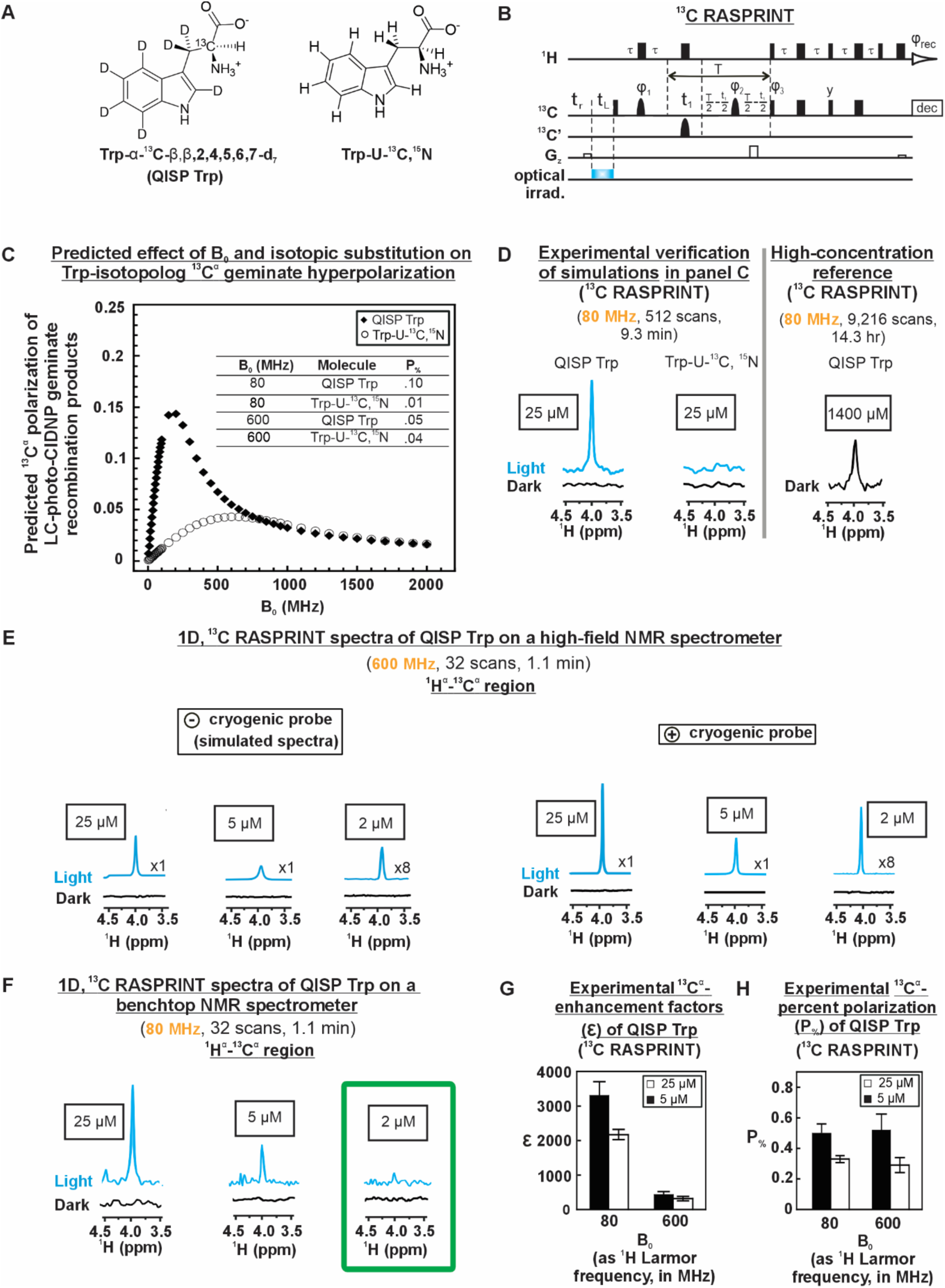
Optically enhanced benchtop NMR spectroscopy enables data collection at low-micromolar concentration. **A**. Structures of QISP Trp and Trp-U-^13^C, ^15^N isotopologs. **B**. ^13^C RASPRINT LC-photo-CIDNP pulse sequence used for this work, tailored for the detection of ^13^C^α^-^1^H^α^. Bell-shaped curves denote 180° Q3_surbop shaped pulses. The parameters tr, tL and T denote the recycle delay, LED irradiation time and total evolution time in the indirect dimension (set to 26.6 ms for C^α^), respectively, and ι− = 1/(4Js_CH_). The phase cycling is *φ*_rec_ is x, −x, −x, x; *φ*, = y, −y; *φ*_2_ = y, y, −x, −x, −y, −y, x, x; *φ*_3_ = x.^31,35^ **C**. Predicted geminate polarization of QISP Trp and Trp-U-^13^C, ^15^N isotopologs at the ^13^C^α^ position as a function of applied magnetic field (B0, in MHz). Simulations were performed with the known g-factors of Trp cation radical, fluorescein anion radical and the corresponding hyperfine coupling constants (see **Supplementary Information**). **D**. ^13^C RASPRINT experiments (under light and dark conditions, left) on 25 μM QISP Trp and Trp-U-^13^C, ^15^N and an only dark spectrum (LED-off) of 1.4 mM QISP Trp at 80 MHz was shown (right) for reference. **E**. ^13^C RASPRINT experiments (under light and dark conditions) on 25 μM, 5 μM and 2 μM QISP Trp at 600 MHz without (left, simulated) and with (right) the cryoprobe. The vertical scale was increased 8-fold for 2 µM QISP Trp at 600 MHz, for better visualization. **F**. ^13^C RASPRINT experiments (under light and dark conditions) on the benchtop NMR on 25 μM, 5 μM and 2 μM QISP Trp. **G**. The bar graphs illustrating the LC-photo-CIDNP enhancement factors (**ε**) for 25 μM and 5 μM QISP Trp with ^13^C RASPRINT at 600 and 80 MHz. **H**. The bar graph illustrating the percent polarization (P%) for 25 μM and 5 μM QISP Trp at 600 MHz and 80 MHz. **ε** and P% were calculated with 32-scan spectra. ^13^C RASPRINT LC-photo-CIDNP spectra shown here were in 10 mM phosphate buffer (pH ∼7.2), used a 0.5 s recycle delay, and 0.2 s (Fig. 2D, left) or 1 s (Figure D-H) LED irradiation time per scan. 25 μM and 5 μM QISP Trp used 12 μM and 8 μM fluorescein as the dye, respectively. Spectra enclosed within a green frame pertain to the lowest concentration detected in this class of experiments. All data are shown as average ± SE (n = 2).

The computational predictions of **Fig. 2C** provide an excellent platform for experimental verification. QISP Trp was synthesized in-house as described,^43^ and ^1^H-detected ¹³C RASPRINT experiments were run (**Fig. 2D**). A short (0.2 s) *in situ* ¹³C^α^ hyperpolarization buildup time with optical irradiation at 453 nm was employed, followed by magnetization transfer to the highly sensitive ¹H for selective detection of the ¹³C^α^–¹H^α^ pair via the ^13^C RASPRINT pulse sequence.^31^ At 25 μM concentration, QISP Trp shows excellent S/N while Trp-U-¹³C, ¹⁵N at 25 µM displays no detectable resonance under light conditions (LED-on,).

In summary the experimental data on the two Trp isotopologs are consistent with theoretical predictions, and QISP Trp gets much more strongly hyperpolarized than Trp-U-¹³C, ¹⁵N at 25 μM concentration, on a benchtop NMR spectrometer at low field (80 MHz, 1.88 T).

Notably, the experiment on QISP Trp also provides a significant advantage in terms of data collection time. This favorable outcome (**Fig. 2D**) is contributed by the short experimental time of the ^13^C RASPRINT experiment, which can employ fairly short recycle delays.^31^ This rapid-data-collection advantage is qualitatively testified by the control dark (LED-off) experiment of **Fig. 2D**, run on a much more concentrated QISP Trp sample with many more scans. For this experiment, the S/N is only half of the value obtained in the light experiment. Considering differences in concentration and number of scans, it would have taken 12,544 times longer than the light-conditions experiment, to gain the same S/N on a 25 μM sample under dark conditions. Therefore, the savings in data-collection time on the benchtop spectrometer, for QISP Trp by LC-photo-CIDNP, is dramatic.

### QISP Trp is readily detected at 2 μM concentration on a benchtop NMR spectrometer

Next, we probed the detectability limit of the QISP Trp isotopolog by directly comparing outcomes by benchtop (80 MHz, 1.88 T) and high-field (600 MHz, 14.1 T) NMR at the lowest possible concentrations granting detectability. The results, reported in **Fig. 2E-F**, show that 2 μM Trp is readily detected at 600 MHz in the presence of a cryogenic probe. In addition, the simulated spectrum of the same sample at 600 MHz in the absence of a cryogenic probe (upon taking standard relative performances into account^32^) also displays a strong predicted signal-to-noise (S/N) (see **Supplementary Information** for details). Similar results are obtained at 80 MHz on a benchtop NMR spectrometer with a room-temperature probe, though with a more moderate S/N. Remarkably, 2 μM QISP Trp data collection on a benchtop NMR spectrometer only takes 1.1 min, with a carefully optimized LED irradiation time of 1 s/scan (**Supplementary Fig. 1A**). This short time, coupled with high sensitivity, renders the combination of benchtop NMR and LC-photo-CIDNP extremely advantageous and desirable. The fluorescein photosensitizer was key for a successful outcome, due to the superior performance of this dye for low-concentration samples, arising from its long photoexcited triplet-state lifetime.^21^

A more quantitative assessment of the advantages provided by LC-photo-CIDNP on benchtop NMR spectrometers (80 MHz, 1.88 T) relative to high field instruments (600 MHz, 14.1 T) was achieved by evaluating the LC-photo-CIDNP enhancement factors (**ε**), defined as

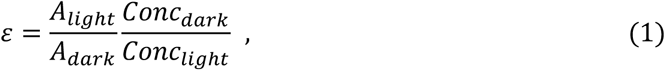

where A and Conc denote resonance areas and sample concentrations, respectively. Note that experiments under dark (LED-off) conditions had to be carried out at much higher concentration than light (LED-on) experiments, to compute χ values. As shown in **Fig. 2G**, ε values at 80 MHz are substantially larger than those at 600 MHz. Namely, at 80 MHz, the χ values of 25 μM and 5 μM QISP Trp were 2,183 ± 102 and 3,100 ± 212, respectively. The corresponding values at 600 MHz were only 243 ± 29 and 432 ± 61, respectively (see also **Supplementary Table 3**). To the best of our knowledge, the 3,100 ± 212 χ of 5 µM QISP Trp is the highest enhancement value reported on a benchtop system to date. Note that higher χ values were only obtained by SABRE for N-containing model compounds at much higher sample concentrations than in the present report (**Supplementary Table 1**).

LC-photo-CIDNP signal intensities exhibit an unconventional dependence on sample concentration, yielding significantly higher ε values at lower sample concentration.^21,22^ This favorable outcome is likely a consequence of the fact that the optimized fluorescein-dye concentration of the 5 μM samples is lower than the optimized dye concentration of the 25 μM samples. As a consequence, fluorescein achieves a higher steady-state population of photoexcited triplet state at lower concentration, given the smaller extent of unproductive self-quenching arising from bimolecular collisions of the photoexcited dye.^21^ Thus, as can be noted in **Fig. 2G** upon comparing χ values at 5 μM and 25 μM, χ values are higher at lower sample concentration. As expected, this relative behavior applies both at low and high applied field B_0_. This type of effect was already experimentally observed and computationally predicted in the past.^21^

The ¹³C^α^ T₁ values of QISP Trp at 80 MHz (T_1, Cα_ = 1.37 ± 0.14 s, determined in this work) and 600 MHz (T_1, Cα_ = 1.56 ± 0.02 s)^18^ are all identical within error (see **Supplementary Information** for details). Therefore, we can rule out any potential contributions to the light effects arising from differential nuclear ^13^C T₁ values, under the steady-state irradiation conditions of this experiment.^27,44^ Moreover, the dark effects associated with QISP isotopologs due to ¹H^α^ linewidth reduction and elimination of coherence losses during the pulse sequence are expected to be B_0_-independent for small molecules, hence essentially equal at 80 and 600 MHz.^33,19^ Therefore, these dark effects are not responsible for the increased enhancements χ observed at low B_0_ in **Fig. 2G**. Differential B_0_-dependent contributions to the final LC-photo-CIDNP spectra due to F-pair polarization are governed by similar principles as those underlying geminate polarization, given that cancellation and degenerate electron exchange effects are likely small, under the low-concentration conditions of our experiments.^33,19^

The corresponding percent polarization (*P*_%_) values shown in **Fig. 2H** account for both Boltzmann distributions at different B_0_ and **χ**. The *P*_%_ parameter is defined according to relations 2 and 3 below

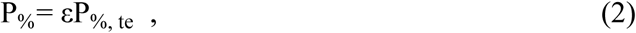

where, *P*_%,te_ is thermal polarization at thermal equilibrium. In addition,

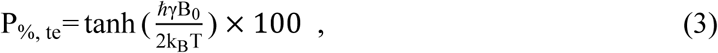

where, ℏ is the reduced Planck’s constant (h/2π), *γ* is the gyromagnetic ratio of the nucleus of interest, *k_B_* is the Boltzmann constant and *T* is the absolute temperature. At 80 MHz, *P*_%_ of 25 μM and 5 μM QISP Trp are 0.37 % ± 0.02 and 0.50 % ± 0.03, respectively, and the corresponding values at 600 MHz are 0.29 % ± 0.03 and 0.52 % ± 0.07 respectively. Remarkably, the identical *P*_%_ values (within error) at the two fields show that LC-photo-CIDNP effectively eliminates the polarization gap between the 80 and 600 MHz applied fields (**Supplementary Fig. 2**).

Overall, the data in **Fig. 2E-H** show that LC-photo-CIDNP renders benchtop NMR spectroscopy (80 MHz) of comparable performance to high-field (600 MHz) NMR. This advantage, combined with results on natural-abundance isotopologs in the sections below, leads the way to *in situ* low-cost bioanalytical NMR spectroscopy of aromatic compounds.

### Natural-isotopic-abundance aromatic compounds can be readily detected at low-μM concentrations by optically enhanced benchtop NMR

We explored hyperpolarization-enhanced benchtop NMR on molecules at natural isotopic abundance, to generalize the sensitivity advantage to more readily available starting materials than QISP amino acids. Towards this end, we computationally assessed the predicted geminate polarization of all protons of unlabeled Trp (**Fig. 3B**). Surprisingly, we identified a strong B_0_ dependence of LC-photo-CIDNP geminate polarization, with favorably larger values at low field, as shown in **Fig. 3B**. For instance, the predicted unlabeled Trp ^1^H^β^ geminate polarization maximum is achieved at ca. 4.7 T (200 MHz). We then proceeded to experimentally test and optimize this label-free approach. To most conveniently perform ^1^H-photo-CIDNP in protonated buffer, we coupled WET (Water Suppression Enhanced through T₁) solvent suppression^45^ with ¹H-photo-CIDNP and developed the ¹H <u>P</u>ulse-<u>A</u>cquire <u>S</u>olvent <u>S</u>uppression via <u>WET</u> (PASS-WET) LC-photo-CIDNP sequence **(Fig. 3C**). Solvent suppression is particularly challenging at low concentration and low B_0_, where technical challenges are exacerbated by the reduced chemical shift dispersion and sensitivity. Here, WET was selected over other solvent-suppression schemes because of its narrower suppression bandwidth, which enables efficient resonance detection even near the residual HDO signal (**Supplementary Fig. 3**). In ¹H PASS-WET LC-photo-CIDNP, for optimal results the LED is kept on throughout the WET selective pulses and turned off just before the first 90° pulse. As a reference, the ^1^H PASS-W5ES LC-photo-CIDNP pulse sequence (**Fig. 3D**) was used to generate data at 600 MHz (**Fig. 3E**) in the absence and presence of a cryogenic probe. Benchtop NMR (80 MHz) data acquired with ^1^H PASS-WET LC-photo-CIDNP are shown in **Fig. 3F**. Conveniently, 5 μM Trp was readily detected in just ∼ 2 min (32 scans) with an optimized 1 s LED irradiation per scan (**Supplementary Fig. 1B**) in aqueous buffer. In contrast, a standard pulse-acquire experiment under dark conditions is estimated to require over 1900 days, to reach a comparable signal-to-noise ratio.

**Figure 3.**
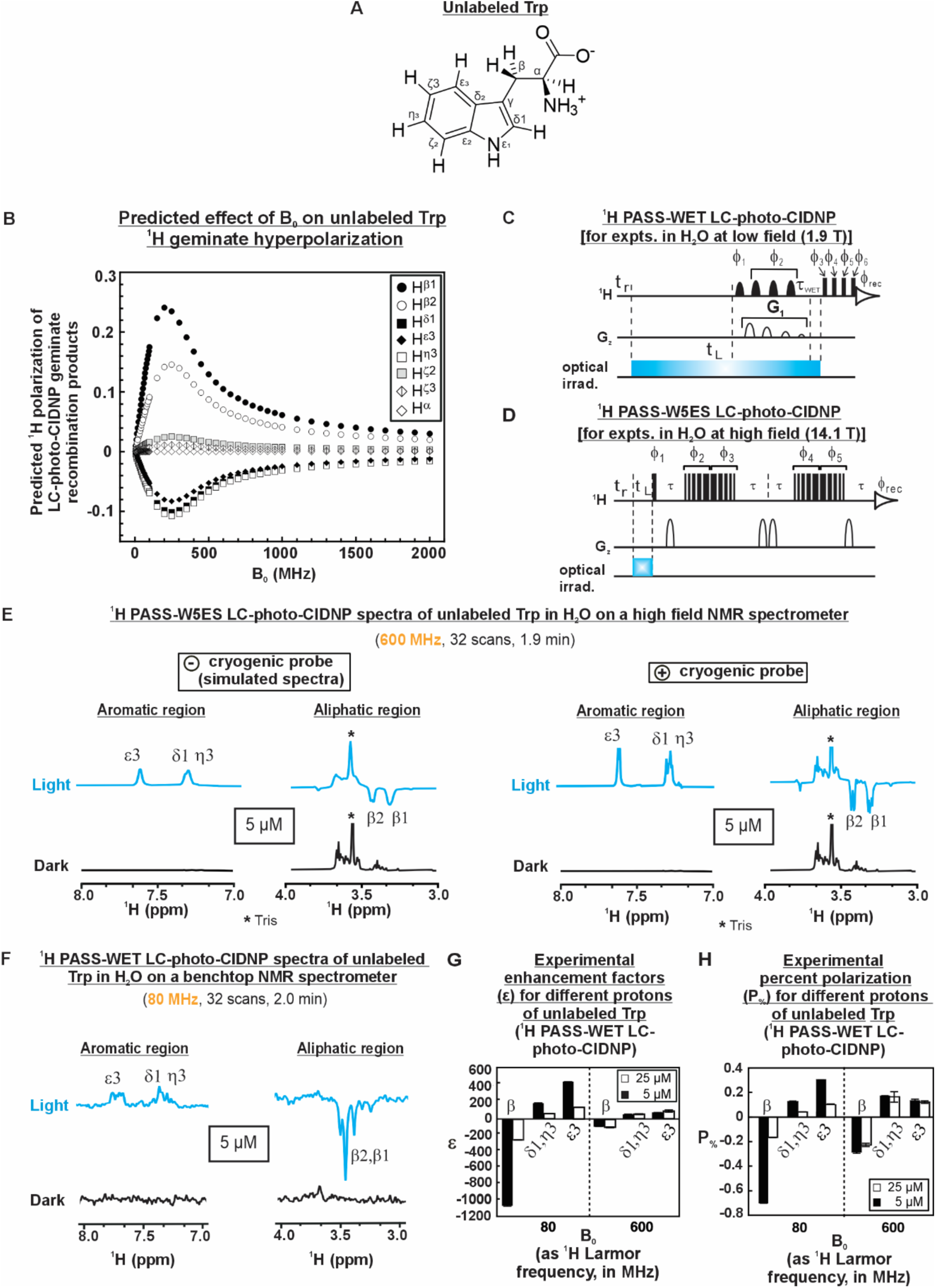
Low-micromolar concentrations of unlabeled molecules of interest are detected via ^1^H PASS-WET LC-photo-CIDNP in aqueous buffer via benchtop NMR. **A.** Structure of unlabeled Trp, **B**. Computationally predicted ^1^H-geminate polarizations of different protons of unlabeled Trp as a function of applied magnetic field (B0, in MHz). Simulations were performed with the known g-factors of Trp cation radical and fluorescein anion radical, and the corresponding hyperfine coupling constants (see **Supplementary Information**). **C**. ^1^H PASS-WET LC-photo-CIDNP pulse sequence. The parameters tr, tL, ι−WET denote recycle delay, LED irradiation time and WET delay period, respectively. ι−WET is (10 ms − ι−) and ι− is 2 ms. SMSQ10.100 gradients (G1) were used at 80%, 40%, 20% and 10%, respectively. *φ_rec_* is x, −x, x, −x, y, −y, y, −y; *φ*_1_ = x; *φ*_2_= y, *φ*_3_ = x, −y, −y, y, −x, x, x, −x, *φ*_4_ =-x, x, −x, x, −y, y, −y, y; *φ*_5_ = −y, y, y, −y, x, −x, −x, x; *φ*_6_ = x, −x, x, −x, y, −y, y, −y. **D**. ^1^H PASS-W5ES LC-photo-CIDNP pulse sequence. The phase cycling is *φ_rec_* is x, −x, x, −x, y, −y, y, −y, *φ*_1_ = x; *φ*_2_= y, *φ*_3_ = x, y, y, x, −x, x, x, −x; *φ*_4_ =-x, x, −x, x, −y, y, −y, y; *φ*_5_ = −y, y, y, −y, x, −x, −x, x. **E**. ^1^H PASS-W5ES LC-photo-CIDNP at 600 MHz on a 5 μM unlabeled Trp (n = 2) without (left, simulated) and with (right) the cryoprobe (under light and dark conditions). **F**. ^1^H PASS-WET LC-photo-CIDNP experiments on the benchtop NMR on a 5 μM unlabeled Trp (under light and dark conditions). **G**. The bar graphs displaying the LC-photo-CIDNP enhancement factors (**ε**, left) and **H**. percent polarization (P%, right). All LC-photo-CIDNP spectra shown here were in 10 mM phosphate buffer (pH ∼7.2), used a 1 s recycle delay and 1 s LED irradiation time per scan. 25 μM and 5 μM unlabeled Trp used 12 μM and 8 μM fluorescein as the dye, respectively. All experiments shown here employed glucose-d₁₂ to minimize background from (regular protonated glucose (glucose-h₁₂) in the aliphatic region (**Supplementary** Fig. 15). Durations of LED irradiation times in panels C and D are not drawn to scale, relative to the actual experimental timings. All data are shown as average ± SE (n = 2).

We then proceeded to determine the hyperpolarization enhancement factor **ε** according to equation 1. This impressive experimental trend is consistent with the theoretical predictions of **Fig. 3A**. Specifically, the **ε** values at 80 MHz are: −1,080.8 ± 4.0 for ^1^H^β^, 194.6 ± 5.2 for ^1^H^δ1^ and ^1^H^ρι3^, 473.3 ± 1.1 for ^1^H^ε3^ at 5 µM (**Supplementary Table 4**). In contrast, the ε values at 600 MHz are much lower: −45.6 ± 11.3 for ^1^H^β^, 35.7 ± 6.8 for ^1^H^δ1^ and ^1^H^ρι3^, 27.4 ± 0.6 for ^1^H^ε3^ at (**Supplementary Table 5)** the same sample concentration. Remarkably, the unlabeled Trp **ε** values at 80 MHz exceed those at 600 MHz for all the hyperpolarized protons (**Fig. 3G**). The similar ^1^H T₁ values at both fields (**Supplementary Table 6**) rule out any potential field-dependent contributions from differences in longitudinal relaxation. Although these ^1^H T₁ values were measured in a D₂O buffer, the field-dependent trends are expected to be similar in aqueous buffer. As in the case of the ¹³C RASPRINT investigations discussed in the previous section, lowering sample concentration to 5 μM leads to larger **ε** and P_%_ values at 80 MHz relative to 600 MHz **(Fig. 3 G, H)**. This effect is likely due to the same dye-concentration-related arguments discussed in the previous section.

In all, the extremely large **ε** and P% values for the aliphatic and aromatic resonance of natural-abundance Trp at 80 MHz expand the practical applicability of benchtop NMR to very-low-concentration aromatic-containing molecules.

### LC-photo-CIDNP on a benchtop NMR spectrometer enables detection of neurotransmitters and pharmaceuticals at nM levels

We extended ¹H PASS-WET LC-photo-CIDNP to a variety of unlabeled small medically relevant compounds, upon taking advantage of their favorable redox properties. We focused on a variety of neurotransmitters that happen to also be downstream metabolites of Trp (serotonin, melatonin) and Tyr (epinephrine and L-DOPA), two classical LC-photo-CIDNP substrates. In addition, we analyzed a variety of well-known small-molecule pharmaceuticals, including a uremic toxin (indoxyl sulfate), two migraine treating pharmaceuticals (zolmitriptan and rizatriptan), a monoamine oxidase inhibitor and potential probe for Parkinson’s (harmaline) and an antihypertensive compound (reserpine). All of the above targets bear one or more aromatic rings and are often the subject of biomedical studies investigating metabolic fate, bioactivity, dosing for pharmacokinetic purposes, and therapeutic monitoring.^46,47^ Low-field NMR for the direct detection at low-μM levels of these compounds has not yet been feasible, under physiologically relevant conditions.

^1^H PASS-WET enabled the ultra-rapid detection of Tyr, epinephrine and serotonin at 1 µM – 500 nM levels (**Fig. 4B,C,F**). An epinephrine photoproduct^33^ was also detected at 250 nM (**Supplementary Fig. 4**). To the best of our knowledge, these are the lowest concentrations ever detected by benchtop NMR in protonated buffer. Further, L-DOPA and melatonin were revealed at 5 µM **(Fig. 4A,E**), while indoxyl sulfate required 25 µM **(Fig. 4D)** levels. Finally, the rizatriptan, zolmitriptan, harmaline and reserpine pharmaceuticals (**Fig. 4G–J**) were detected at 25 – 100 µM concentrations. See sections below for the biomedical relevance of the above concentrations. **Supplementary Fig. 5,6** reports the chemical shift assignments. The fluorescein photosensitizer was used for all the indole-like compounds, while ATTO Thio 12 was employed for all the hydroxy-benzene-like molecules.^21,48^ It is worth noting that, the compounds with electron rich aromatic rings are expected to perform well in LC-photo-CIDNP hyperpolarization due to their high electron density and low redox potentials, which facilitates the transient triplet-state radical-pair formation, which is a prerequisite of this technology.^49^ In the case of harmaline and reserpine, trace amounts of DMSO improved the signal-to-noise ratio **(Supplementary Fig. 7)**, likely due to solubility, viscosity or redox-potential effects (see **Supplementary Information**).^50^ The exact role of DMSO, however, remains unknown, and beyond the scope of this work.

**Figure 4.**
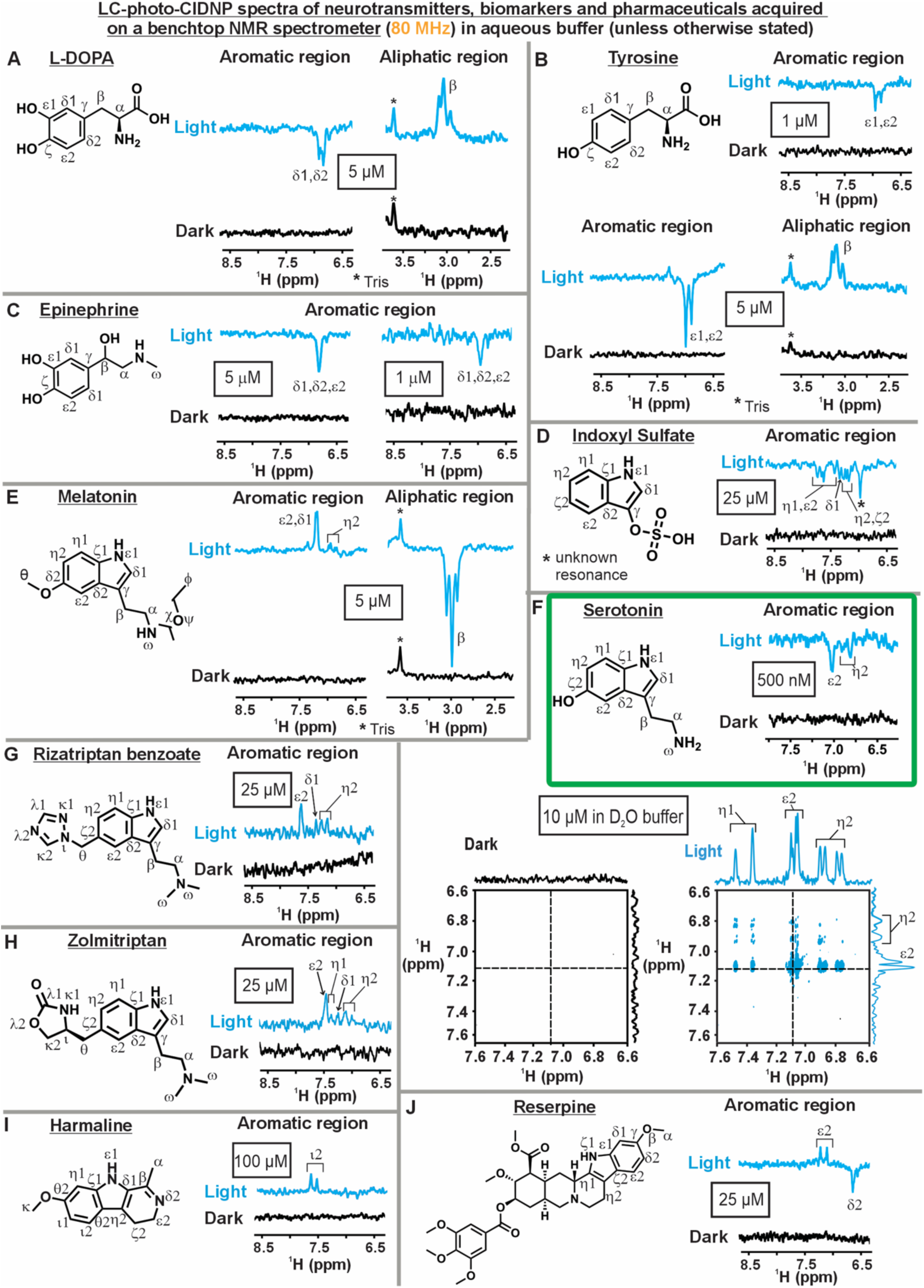
Benchtop 1D/2D LC-photo-CIDNP NMR enables low-micromolar detection of biomedically relevant compounds. LC-photo-CIDNP spectra (under light and dark conditions) of **A**. L-DOPA at 5 µM (64 scans, 3.5 min), **B**. Tyrosine at 1 µM (64 scans, 3.5 min, top) and 5 μM (64 scans, 3.5 min, bottom). **C**. Epinephrine at 5 μM (16 scans, 0.9 min) and 1 μM (88 scans, 4.8 min). **D**. Indoxyl sulfate at 25 μM (256 scans, 14.0 min). **E**. Melatonin at 5 μM (128 scans, 7.0 min). **F**. Serotonin in H2O at 500 nM (top, 1D, 128 scans, 7.0 min) and in D2O at 10 μM (bottom, 2D, 4 scans/transient, 128 transients, 22.0 min) **G**. Rizatriptan benzoate at 25 μM (16 scans, 0.9 min). Horizontal and vertical black dashed lines in the 2D spectra indicate the corresponding 1D ^1^H projections, shown along the respective axes in black (dark) and blue (light), **H.** Zolmitriptan at 25 μM (16 scans, 0.9 min). **I.** Harmaline at 100 μM (32 scans, 1.8 min) and **J.** Reserpine at 25 μM (128 scans, 7.0 min). Extra amounts of DMSO-d6 were added to harmaline and reserpine (see **Supplementary Information** and **Supplementary** Fig. 7) for improved performance. All spectra shown here (except for 2D experiments) were acquired with the ^1^H PASS-WET pulse sequence (shown in Fig. 3C) and in 100% aqueous buffer (potassium phosphate, pH ∼7.2). The 2D data were collected with the COSY-LC-photo-CIDNP pulse sequence (shown in **Supplementary** Fig. 8) in 100% D2O buffer (potassium phosphate, pH ∼7.2). All LC-photo-CIDNP experiments employed a 1 s recycle delay and 1 s LED (A–J, except 2D) irradiation time/scan, while the 2D data employed 0.2 s LED irradiation/scan. All 1 μM and 5 μM phenolic scaffolds used 5 μM and 10 μM ATTO Thio 12, respectively. All 1 μM, 5 μM, 25 μM and 100 μM indole-scaffolds used 2.5 μM, 8 μM, 12 μM and 25 μM fluorescein, respectively. 2D LC-photo-CIDNP experiment with 10 μM serotonin used 15 μM fluorescein. All experiments shown here were performed with n = 2. See **Supplementary** Fig. 5**,6** for chemical shift assignments. All experiments shown here employed glucose-d₁₂ to minimize background signal from regular protonated glucose (glucose-h₁₂) in the aliphatic region (**Supplementary** Fig. 15). Data enclosed within a green frame highlight the lowest concentration detected in this class of experiments. All data were collected on a 1.88 T (80 MHz) benchtop NMR spectrometer.

### Two-dimensional benchtop LC-photo-CIDNP improves spectral resolution and enables hyperpolarization transfer

While ^1^H PASS-WET LC-photo-CIDNP significantly enhances sensitivity, resolution is limited on an 80 MHz magnet and few resonances get hyperpolarized. To overcome these limitations, we implemented the 2D COSY LC-photo-CIDNP pulse sequence (**Supplementary Fig. 8**). Unlike conventional COSY (**Supplementary Fig. 9**), COSY LC-photo-CIDNP spectra are not symmetric about the diagonal. Following frequency labeling during the evolution period (t₁), COSY LC-photo-CIDNP employs a 90° pulse to enable hyperpolarization transfer from photo-CIDNP-active nuclei to their J-coupled partners. Consequently, resonances that are inactive in traditional 1D photo-CIDNP become indirectly hyperpolarized and detectable in the direct (f_2,_ here ^1^H) dimension (see **Supplementary Information** for details). To further improve spectral resolution, we restricted the spectral width to 3.0 ppm, focusing on the aromatic region. **Fig. 4F** shows 2D COSY LC-photo-CIDNP spectra on 10 μM serotonin. Data were acquired in ca. 20 minutes. The 1D projection along the direct dimension reveals the ^1^H^η1^ resonance, which was indirectly hyperpolarized *via* ^1^H^ε2^ and ^1^H^η2^, as confirmed by the connectivities between cross and diagonal peaks. This 2D strategy provides high-resolution spectral fingerprints for the accurate identification of aromatic resonances despite the low field of benchtop spectrometers.

### The benchtop-NMR hyperpolarization toolkit developed here leads to hypersensitive nanomolar-level detection of pure neurotransmitters in buffer

Given the encouraging results discussed in the previous section, which focused on the hypersensitive detection of a variety of neurotransmitters and pharmaceuticals, we undertook a more systematic approach to the development of a tailored benchtop-NMR toolkit for samples in aqueous buffer. We focused on three main ^1^H LC-photo-CIDNP benchtop tools: ^1^H PASS-WET, solvent deuteration, and resolution enhancement by homonuclear decoupling via the SHARPER pulse sequence. Each of these tools is discussed below. Serotonin (**Fig. 5A**) was employed as a representative molecule of interest.

**Figure 5.**
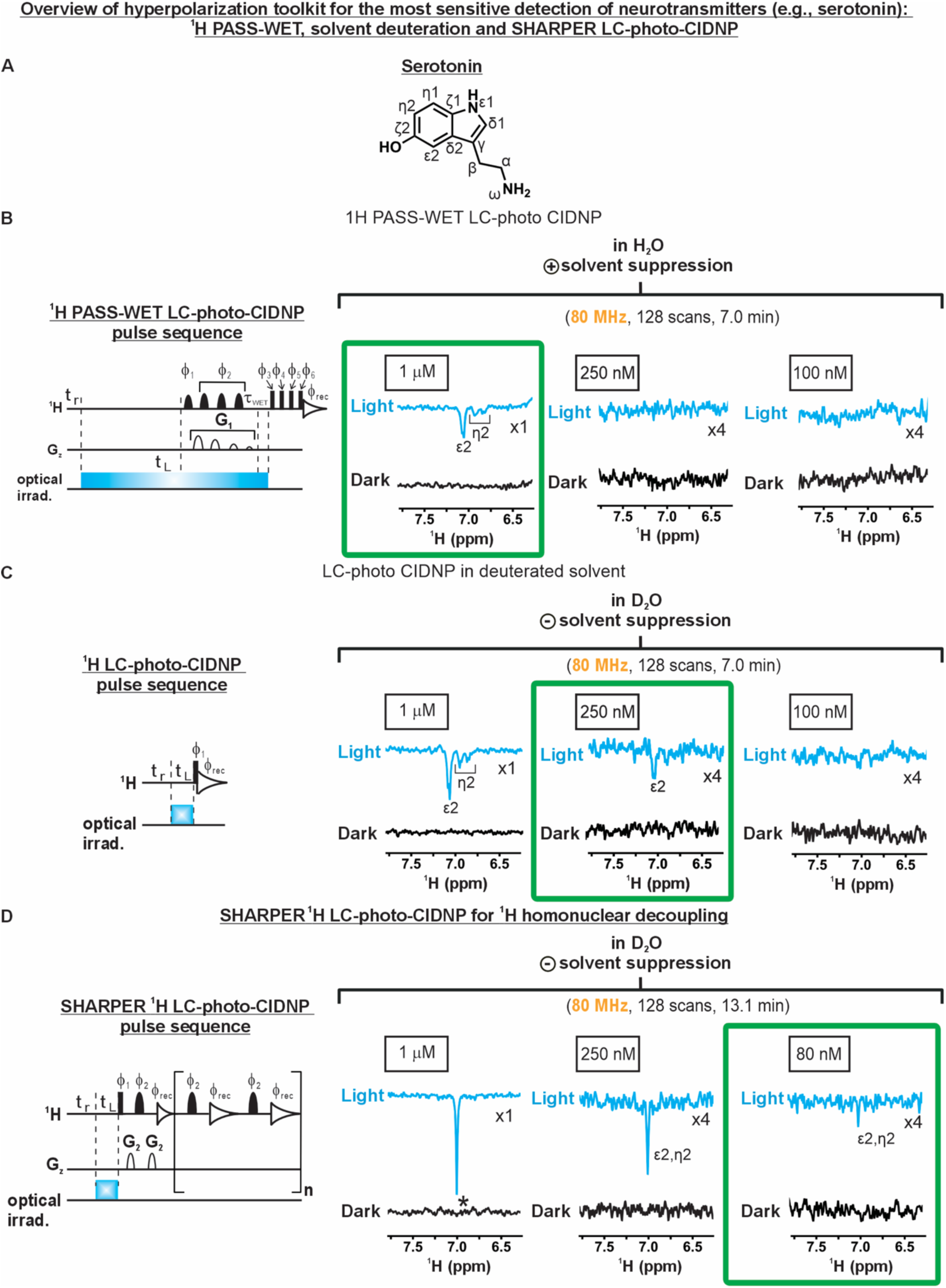
Benchtop 1D ^1^H LC-photo-CIDNP NMR enables detection of biomedically relevant compounds at nanomolar concentration. **A.** Structure of serotonin. **B**. ^1^H PASS-WET LC-photo-CIDNP experiments (pulse sequence on the left) acquired on 1 μM, 250 nM and 100 nM serotonin in 100% aqueous buffer (pH ∼7.2) under light (LED-on) and dark (LED-off) conditions. **C**. ^1^H LC-photo-CIDNP experiments without solvent suppression (pulse sequence on the left) acquired on 1 μM, 250 nM and 100 nM serotonin in 100% D2O-based buffer (pH ∼7.2) under light (LED-on) and dark (LED-off) conditions. **D.** SHARPER ^1^H LC-photo-CIDNP experiments (pulse sequence on the left) acquired on 1 μM, 250 nM and 80 nM serotonin in 100% D2O-based phosphate buffer (pH ∼7.2) under light (LED-on) and dark (LED-off) conditions. Under SHARPER ^1^H LC-photo-CIDNP conditions, 80 nM serotonin is readily detected (green box) in the light (LED-on) spectrum. Dark rectangular features denote nonselective 90° pulses and dark bell-shaped features denote selective 180° refocusing pulses. Here, τ = AQ/2*n*, where AQ is the acquisition time and *n* is the number of loops. G2 denotes an SMSQ10.100 gradient pulse. For SHARPER LC-photo-CIDNP, the following phase cycles were used: β1 = *x*; β2 = y, −y, −x, x; β3 = y, −y; βrec = x, x, −x, −x (see methods for details). The recycle delay and LED irradiation time per scan are denoted by *t*r and *t*L, respectively. All 1 μM serotonin samples contained 2.5 μM fluorescein; all 100 nM, 80 nM and 250 nM serotonin samples contained 500 nM fluorescein (see **Supplementary** Fig. 16). All experiments shown here used glucose-h12. Spectra were acquired using a 1 s recycle delay and 1 s LED irradiation per scan. Each dataset represents at least two independent experiments (*n* ≥ 2). The data enclosed within a green frame highlight the lowest concentration detected in this class of experiments. All data were acquired on a 1.88 T (80 MHz) benchtop NMR spectrometer.

As shown in **Fig. 5B** and as discussed in the previous section, the ^1^H PASS-WET LC-photo-CIDNP pulse sequence is particularly appropriate for the hypersensitive data collection on aromatic neurotransmitters, metabolites and pharmaceuticals in aqueous buffer by benchtop NMR. Concentration-challenging experiments on serotonin in protonated buffer (**Fig. 5B**) show clear detectability at 1 μM levels. In addition, data in **Fig. 4F** illustrate that 500 nM concentration is also readily detected.

In addition, we explored ¹H LC-photo-CIDNP benchtop NMR in 100% D₂O as a strategy to further improve sensitivity. As shown in **Fig. 5C**, the S/N of 1 μM serotonin reproducibly increases by ∼35 ± 9.4 % in D₂O relative to H₂O, and this neurotransmitter is detectable down to 250 nM in D₂O. Although D₂O has previously been used in photo-CIDNP,^51–53^ direct assessments and comparisons between data collected in H_2_O and D_2_O have not been carried out before. The better LC-photo-CIDNP performance in D_2_O is expected to arise from two main factors. First, in D₂O the exchangeable indole N–H proton is replaced by deuterium (at pH ca. 6-7), decreasing the extent of proton-driven dipolar relaxation and extending T₁ relaxation times of nearby protons, consistent with the literature and with our own observations for Trp (**Supplementary Table 7**).^54^ **Supplementary Fig. 10** shows the ^1^H PASS-WET-like inversion-recovery sequence used to measure ^1^H T₁ at 80 MHz. A longer T_1_ is favorable for steady-state photo-CIDNP because it prolongs the lifetime of the hyperpolarized state.^27,44^ Second, the HN indole proton of Trp has a large hyperfine coupling constant (HFC), whose effect on the frequency of triplet singlet mixing is to decrease it, thus lowering the observed LC-photo-CIDNP resonance enhancement.^19,33,52^ Deuteration of the HN indole proton attenuates the signal loss due to the latter effect, thus favoring LC-photo-CIDNP hyperpolarization.

Finally, to enhance both resolution and sensitivity of optically enhanced benchtop NMR, we developed ^1^H SHARPER-LC-photo-CIDNP, which accomplishes homonuclear decoupling during data acquisition. As shown in **Fig. 5D**, using this approach, serotonin can be readily detected down to 80 nM concentration in D₂O. This outcome corresponds to the lowest sample concentration ever detected on a benchtop NMR spectrometer. SHARPER LC-photo-CIDNP employs a series of CPMG-style refocusing π pulses interleaved with brief periods of data acquisition. The complete free induction decay is then reconstructed from the signal portions collected in between successive π pulses (see Methods for details).^55,56^ As a result, this approach not only collapses the selected scalar-coupled multiplets into a single resonance but also suppresses any magnetic field inhomogeneities. Of course, the “spectral sharpening” comes at the expense of J-coupling information. In addition, any distinct resonances that are originally excited collapse into a single signal (chemical-shift collapse). Therefore, chemical shift information may be lost if the excitation pulses are not sufficiently selective to only excite individual resonances. The latter caveat is not usually an issue in LC-photo-CIDNP, whether or not experiments are performed on a benchtop instrument, given that it is typically feasible to employ shaped pulses to excite individual resonances.

### LC-photo-CIDNP on a benchtop NMR spectrometer enables background-free detection of low-µM neurotransmitters and metabolites in prokaryotic and eukaryotic media

We explored the performance of benchtop LC-photo-CIDNP NMR in complex biological environments. We carried out experiments in an *E. coli* cell-like medium, which was prepared in-house based on a modified version of a known protocol,^57^ as outlined in **Fig. 6A,B**. This medium contains most cellular components (e.g., proteins, small molecules, ribosomes, tRNAs) except for genomic DNA and cell membranes, which were eliminated via centrifugation. Remarkably, use of this medium enabled prompt *in-situ* detection of 5 μM QISP Trp via ^1^H-detected ^13^C LC-photo-CIDNP (¹³C RASPRINT), as shown in **Fig. 6C**. In addition, natural-abundance Trp was also detected at 25 μM levels via ¹H PASS-WET LC-photo-CIDNP **(Fig. 6D)**. In the case of ¹³C RASPRINT, given that endogenous cellular components are not ¹³C-enriched and thus fail to produce detectable spectral features, the spectra are much “cleaner” and devoid of resonances not belonging to the molecule of interest. In the case of ¹H PASS-WET LC-photo-CIDNP performed on natural abundance samples, endogenous cellular components are also present in the spectra. On the other hand, these components can be easily identified by control experiments (**Supplementary Fig. 11**), even if some of them may exhibit some degree of LC-photo-CIDNP enhancement. In turn, if desired, endogenous components in ^1^H optically enhanced NMR spectra can be subtracted out by difference spectroscopy.

**Figure 6.**
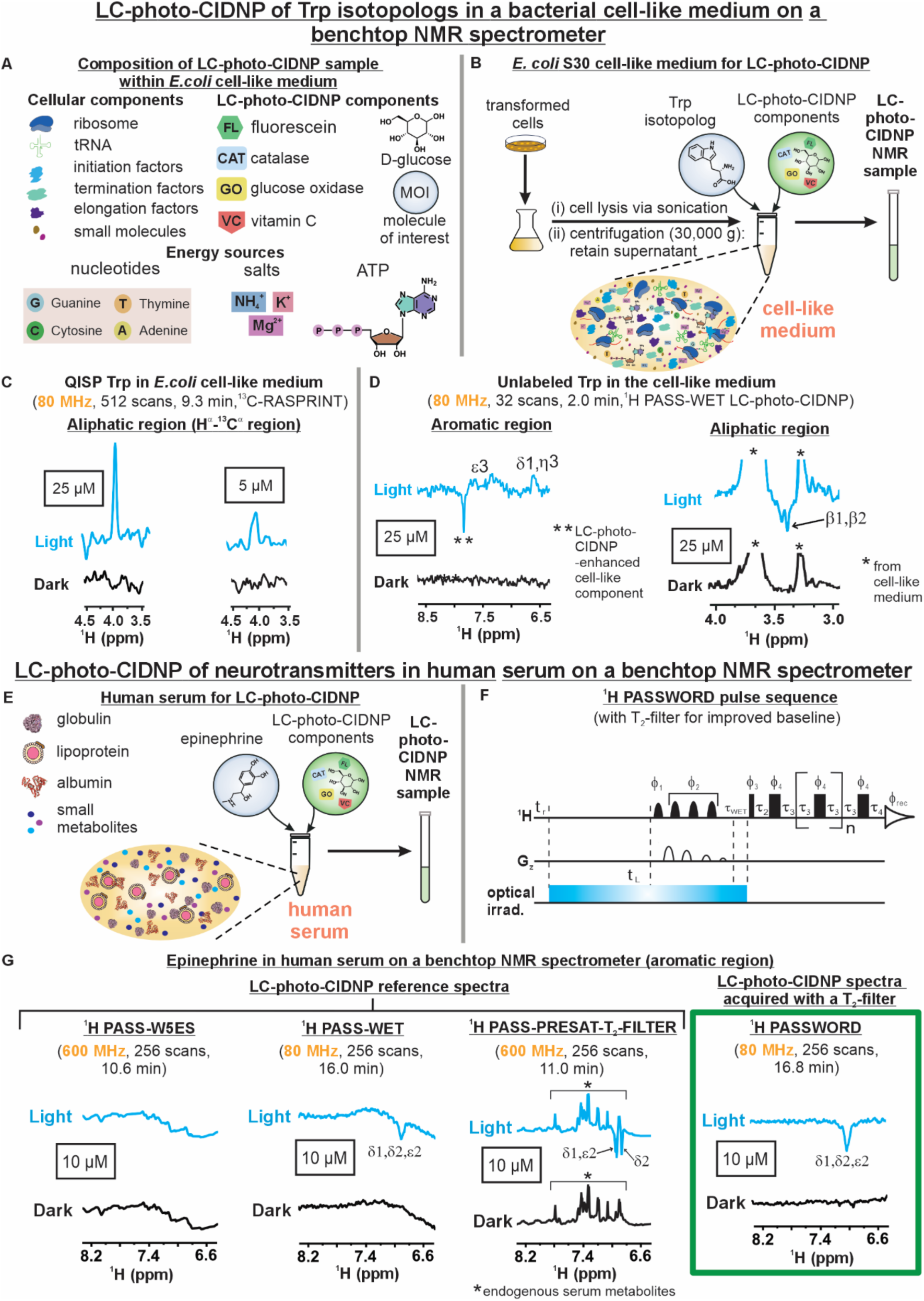
Benchtop LC-photo-CIDNP NMR enables low-micromolar detection of Trp and neurotransmitters in complex prokaryotic and eukaryotic media. **A.** Composition of LC-photo-CIDNP samples including a prokaryotic (bacterial *E. coli*) cell-like medium (see Methods). **B.** Key preparation aspects of *E. coli* cell-like medium for LC-photo-CIDNP experiments (pH ∼ 7.2). **C.** ^13^C RASPRINT spectra of 25 µM and 5 µM QISP Trp (under light and dark conditions, n = 2). **D.** ^1^H PASS-WET LC-photo-CIDNP spectrum of 25 µM unlabeled Trp (under light and dark conditions, n = 2). A set of control experiments were performed to identify the resonances labeled with asterisks (Supplementary Fig. 11). **E**. Composition of LC-photo-CIDNP sample in an unaltered eukaryotic (human serum) medium (pH ∼ 7.1). **F**. ^1^H PASSWORD pulse sequence equipped with T2-filtering and WET solvent suppression (see **Supplementary Information**). **G**. LC-photo-CIDNP spectra of 10 µM epinephrine in human serum. Reference spectra (under light and dark conditions) shown included ^1^H PASS-W5ES at 600 MHz (first, n = 2), ^1^H PASS-WET at 80 MHz (second, n = 2), ^1^H PASS-PRESAT-T2-FILTER at 600 MHz (third, n = 2). Within green box included the spectra (under light and dark conditions, n = 2) with ^1^H PASSWORD at 80 MHz. All LC-photo-CIDNP spectra shown here were prepared in 10 mM phosphate buffer (pH ∼7.2). ^13^C RASPRINT experiments used 0.5 s recycle delay and 0.2 s LED irradiation time per scan. ^1^H PASS-WET experiments in panel D and G both used 1 s recycle delay and 1 s LED irradiation time per scan, respectively. ^1^H PASS-W5ES and ^1^H PASS-PRESAT-T2-FILTER **(Supplementary** Fig. 13) used 1 s recycle delay and 1 s LED irradiation time per scan. ^1^H PASSWORD also used 1 s recycle delay and 1 s LED irradiation time per scan. 25 μM and 5 μM QISP Trp/Trp samples used 12 μM and 8 μM fluorescein as the dye, respectively. 10 μM epinephrine used 10 μM ATTO Thio 12 dye. Data enclosed within a green frame denote the lowest concentration detected in this class of experiments.

Given the clinical relevance of neurotransmitters as disease biomarkers and their particular suitability to display optically driven enhancements, we applied benchtop LC-photo-CIDNP to the analysis of epinephrine, chosen as a representative target, in a eukaryotic matrix, i.e., unmodified human serum.^58,59^ Before the present work (**Fig. 4C**), *in situ* monitoring of epinephrine has been unattainable by NMR due to both this technique’s low sensitivity and the matrix complexity.

Notably, the broad background resonances, due to abundant cellular macromolecules (e.g., globulins, lipoproteins) bearing short spin–spin relaxation times (T_2_), pose a major challenge (**Supplementary Fig. 12A**). This issue can be largely addressed by exploiting the Carr–Purcell–Meiboom–Gill (CPMG) pulse sequence via T_2_ filtering.^60^ The repeated 180° refocusing pulses suppress short-T₂ resonances, enabling clearer detection of low-molecular-weight metabolites, which bear longer T₂s.^61^ Hence, we coupled the advantages of CPMG to ¹H PASS-WET LC-photo-CIDNP and developed ^1^H PASSWORD pulse sequence (<u>P</u>ulse-<u>A</u>cquire with <u>S</u>olvent <u>S</u>uppression via <u>W</u>ET <u>O</u>vercoming <u>R</u>elaxation-related <u>D</u>efects, **Fig. 6F**). ¹H PASSWORD LC-photo-CIDNP yields clean, background-free spectra (**Supplementary Fig. 12B**). We employed this approach to detect epinephrine at 10 µM levels in human serum (**Fig. 6G** green-framed spectrum; serum diluted to 20%) by benchtop NMR in a little over 15 min.

In sharp contrast, when experiments were performed at high field (600 MHz, **Fig. 6G**, first and third reference spectra from the left), the photo-CIDNP-enhanced emissive resonances of epinephrine are heavily obscured by the abundant endogenous serum components. This outcome is due to the much more pronounced sensitivity of the high-field spectrometer under dark conditions. This phenomenon hampers reliable epinephrine detection at 600 MHz, rendering high-field data collection altogether undesirable. Note that at high field T_2_ filtering was accomplished in the presence of solvent suppression via presaturation **(Supplementary Fig. 13),** given the poor performance of WET at high field. On the other hand, aromatic resonances are well separated from the HDO signal (at ca ∼ 4.7 ppm), therefore the solvent-suppression strategy is not expected to affect the outcome.

Conveniently, at low-field (80 MHz) undesirable background resonances are naturally suppressed, given the low sensitivity of the 80 MHz spectrometer under dark conditions. This useful feature enables clear and unambiguous detection of the neurotransmitter LC-photo-CIDNP–enhanced resonances, even in the absence of T_2_-filtering (e.g., **Fig. 6G**, second spectrum from the left), at low field. T_2_-filtering does an excellent job at correcting the rolling baseline in the aromatic region of the 10 μM epinephrine spectrum (**Fig. 6G**, green-framed spectrum). The T_2_-filtering tool also dramatically improves spectral features within the aliphatic region. For instance, it improves the baseline and declutters the spectra of (a) a control sample lacking neurotransmitters (**Supplementary Fig. 12B**) and (b) a corresponding sample including 10 μM epinephrine (**Supplementary Fig. 14**).

In all, the above scenario highlights the fact that strategic exploitation of T_2_ filtering in combination with the low sensitivity of benchtop NMR under dark conditions is highly beneficial for the hypersensitive detection of aromatic neurotransmitters like epinephrine on a benchtop NMR spectrometer. Importantly, neurotransmitters are frequently present at low- to mid-μM concentration in eukaryotic (serum, cerebrospinal fluid) and prokaryotic (bacterial cytoplasm and periplasm) fluids.^2,62^ Moreover, the concentrations of these biomolecules serve as sensitive indicators of the presence of physiological *vs* pathological states.^2,62^ Low micromolar levels match the concentrations readily detected by the optically enhanced benchtop approach reported here, underscoring its suitability for the rapid *in situ* analysis of aromatic-moiety-containing small molecules within complex biological *milieux*.

## Discussion

Understanding the role of biologically relevant molecules including amino acids, neurotransmitters, metabolites, and pharmaceuticals under physiologically relevant conditions remains a central goal in both basic research and clinical/diagnostic practice. Solution-state NMR spectroscopy at low field is uniquely suited to accomplish this task. Unlike other methodologies (e.g., LC-MS, surface-enhanced light scattering, near-infrared fluorescence), benchtop NMR is nonperturbative and provides structural and dynamical information *in situ* at atomic resolution.

In this work, we developed a novel LC-photo-CIDNP toolkit including experimental strategies and NMR pulse sequences to collectively address the main limitations of low-field NMR i.e., sensitivity, spectral resolution, and spectral congestion. By this approach, we enabled facile, high-quality and ultra-rapid analysis of biologically and clinically relevant biomolecules.

It is well-known that catecholamine- and indoleamine-based neurotransmitters are inherently prone to oxidative modification in the brain.^39^ They are also involved in endogenous RP formation via one-electron oxidation.^39^ Conveniently, all neurotransmitters bear one or more electron-donating substituents on their aromatic ring, which donate electron density into the aromatic π-system, stabilizing the radical (cationic or neutral) within photochemically generated radical pairs. These molecular features provide a mechanistic basis for the strong LC-photo-CIDNP responses displayed by neurotransmitters.

The experimental benchtop-NMR strategies introduced here are in principle also applicable to other applied fields, though some of them are specifically tailored to low field (e.g., see **Fig. 6G**). First, we developed ¹H LC-photo-CIDNP with WET solvent suppression (¹H PASS-WET, **Fig. 3D**) for measurements in 100% aqueous buffer with detection of bioanalytes down to 500 nM within minutes **(Fig. 4)**. Second, we combined ¹H PASS-WET with a CPMG T₂ filter (¹H PASSWORD, **Fig. 6F**) to correct baseline distortions from broad linewidths in complex biofluids (e.g., urine and serum). This is a longstanding limitation exacerbated at low field by the reduced chemical shift dispersion.^60^ Using ¹H PASSWORD, epinephrine was detected at 10 μM **(Fig. 6G)** in unmodified human serum in ∼16.8 min without any interfering endogenous resonances from serum. Third, we developed SHARPER ¹H LC-photo-CIDNP (**Fig. 5D**) to enhance sensitivity by collapsing selected resonances into sharp singlets and via eliminating magnetic field inhomogeneity.^55,56^ This strategy enabled detection of 80 nM serotonin (11.45 ng) at natural abundance in 100% D₂O buffer (**Fig. 5D**). This concentration is of significant interest for clinical purposes where dysregulated or elevated plasma or serum serotonin levels are implicated in several diseases including chronic kidney disease (>113 nM serotonin) and carcinoid heart disease (up to 9.75 μM serotonin).^63,64^ In summary, our method achieves sensitivities approaching, though not yet matching, other established bioanalytical techniques including radioenzymatic assays, HPLC-ECD, LC-MS, and ELISA.^64^ On the other hand, none of the above established technologies are able to detect conformational changes or environmental variations *in situ* at atomic resolution, which is a prominent advantage of the optically enhanced benchtop NMR approach introduced here, especially in the case of neurotransmitter detection.

In addition to developing new pulse sequences, we evaluated several existing LC-photo-CIDNP NMR strategies and established their best performances so far on a benchtop spectrometer. Adrian’s radical-pair theory was employed here to predict LC-photo-CIDNP enhancements (**Fig. 2C, 3B**).^18,19,33,40^ We also implemented ¹H-detected ¹³C RASPRINT for the detection ^1^H^α^–^13^C^α^ resonance of selectively deuterated Trp (a.k.a. QISP Trp) at 2 μM in ∼1.1 min **(Fig. 2F)**. Further, these QISP amino acid isotopologs can be easily incorporated into proteins and given the compatibility of LC-photo-CIDNP with macromolecules,^29,31,32,65^ we also envision benchtop LC-photo-CIDNP soon being used for high-resolution protein structural studies. These results highlight deuteration as a strategy for improving LC-photo-CIDNP sensitivity for any molecule of interest. It readily extends beyond ¹³C-photo-CIDNP and we have successfully adapted it to ¹H and ¹⁹F LC-photo-CIDNP (data not shown). We addressed the poor resolution of low-field NMR using 2D COSY LC-photo-CIDNP with a reduced sweep width. This experiment has an added benefit of enabling the detection of photo-CIDNP-inactive resonances via polarization transfer, as shown for serotonin at 10 μM **(Fig. 4F)**. Data collection only takes 22.0 min, which is exceptionally fast for a high-resolution 2D experiment at 80 MHz.

In summary, our results show that benchtop NMR, which employs a magnet ∼7.5-fold smaller than a 600 MHz spectrometer and a room-temperature probe, can achieve a performance comparable to high-field instruments with cryogenic probes. By combining tailored pulse sequences with LC-photo-CIDNP approach, we achieved high sensitivity and atomic-level resolution *in situ*, in both buffered solution and prokaryotic as well as eukaryotic environments. Reproducible detection of biologically relevant aromatic metabolites, including neurotransmitters, only takes a few minutes, and can be achieved down to nanomolar concentrations. Notably, the detection of serotonin at 80 nM represents the lowest analyte concentration reported to date on a benchtop NMR spectrometer. Collectively, these approaches demonstrate orders-of-magnitude reductions in experimental time compared to conventional NMR and other routinely used analytical methods for the detection of neurotransmitters and other aromatic metabolites and pharmaceuticals. In all, this work establishes a new benchmark for low-field benchtop NMR spectroscopy and provides a versatile, readily accessible toolkit for exploring metabolism, signaling and disease diagnostics at a fraction of the cost and complexity of currently employed technologies.

## Acknowledgements

We thank Heike Hofstetter, Gabi Carosio, Clemens Anklin and members of the Cavagnero group for useful discussions. This work was supported by the National Institutes of Health (grant R01GM125995 and R35GM161252 to SC). The Bruker Avance III 600 NMR spectrometer in the Department of Chemistry was supported by NIH grant S10 OD012245.

## Author contributions

S.C., A.H. and S.C.C. designed the research. A.H., S.C.C. and J.H.J. performed the experiments. U.M.S. prepared key materials and media and offered technical advice. C.F.M.C. shared critical hardware expertise and technical assistance that enabled smooth benchtop-spectrometer operation. S.C., A.H., S.C.C. and J.H.J. analyzed the data. A.H., S.C.C., J.H.J. and S.C. wrote the paper.

## Additional information

Supplementary information is available for this paper at https://doi.org/xxx

Reprints and permissions information is available at [TBD URL]. Correspondence and requests for materials should be addressed to S.C.

## Experimental Section

### Materials

Natural-abundance tryptophan (Trp) was purchased from Advanced ChemTech. Uniformly ^13^C, ^15^N labeled Trp (Trp-U-^13^C, ^15^N) was obtained from Cambridge Isotopes Laboratories, Inc. The Trp-α-^13^C-β-β,2,4,5,6,7-d_7_ or QISP Trp was synthesized in house (see below) from formaldehyde-^2^H_2_ and glycine(2-^13^C), which were purchased from Cambridge Isotopes Laboratories, Inc, indole-^2^H_7_ purchased from CDN Isotopes, Pyridoxal-5’-monophosphate (PLP) purchased from MilliporeSigma and catalytic amounts of the *Pyrococcus furious* tryptophan synthase b-subunit 2B9 (*Pf*TrpB^2B9^) and *Thermogota maritima* L-threonine aldolase (*Tm* LTA, Enzyme commission classification code, EC 4.1.2.5) synthesized in the lab following the procedure described in the literature.^43^ The sodium salt of the fluorescein photosensitizer dye, vitamin C (ascorbic acid, VC, EC 200-066-2) and the oxygen scavenging enzymes consisting of *Aspergillus niger* glucose oxidase (GO, EC 1.1.3.4), bovine liver catalase (CAT, EC 1.11.1.6) were purchased in the form of freeze-dried powder from MilliporeSigma. ATTO Thio 12 was purchased from ATTO-TEC GmbH. The D-glucose-h_12_ was purchased from ThermoFisher and D-glucose-d_12_ was purchased from Santa Cruz Biotechnology and Cambridge Isotopes Laboratories, Inc. DMSO-d_6_ was purchased from MilliporeSigma. Unlabeled Tyr was purchased from Advanced ChemTech. Serotonin hydrochloride, reserpine, rizatriptan benzoate salt, harmaline, indoxyl sulfate, 3,4-Dihydroxy-L-Phenylalanine (L-DOPA) were purchased from MilliporeSigma, Melatonin was purchased from Mp Biomedicals Inc. Zolmitriptan was purchased from Targetmol Chemicals Inc. Racemic mixture of epinephrine hydrochloride was purchased from Sigma Aldrich. Human serum (heat inactivated from AB clotted whole blood) was purchased from MilliporeSigma.

### Synthesis of QISP Trp

The QISP Trp (IUPAC name: (S)-2-Amino-3-[(2,4,5,6,7-^2^H_5_)-3-indolyl](2-13C,3,3-^2^H_2_)propionic acid) was prepared and purified in the lab, following a one-pot reaction scheme which is a two-step enzymatic cascade biosynthesis starting from glycine(2-^13^C), Formaldehyde-^2^H_2_, indole-^2^H_7_ in the presence of *Pf* TrpB^2B9^ and *Tm* LTA enzymes and PLP cofactor.^43^ The product was identified, and the overall yield was determined, by ^1^H-NMR. Product identity was also confirmed by electrospray ionization mass spectrometry (ESI-MS).

### Oxygen scavenging enzymes

GO and CAT in presence of D-glucose-h_12_ or −d_12_ were used as the oxygen scavenging enzymes. Powders of GO and CAT were dissolved in a 10 mM H_2_O- or − D_2_O based potassium phosphate buffer (pH 7.2) solution. The concentration of each enzyme was measured by an electronic absorption spectrophotometer with an extinction coefficient of 267,200 M^−1^cm^−1^ at 280 nm for GO and 912,500 M^−1^cm^−1^ at 276 nm for CAT.^66^ Small aliquots of the enzyme stock solutions were made and these were flash-frozen with liquid nitrogen and stored at −80 °C. The frozen aliquots were thawed in a water bath around 4 °C on the day of the experiment. The remaining thawed enzymes were discarded after the experiments.

### Fluorescein and ATTO Thio 12 dye

The sodium salt of the fluorescein dye or ATTO Thio 12 were dissolved in distilled deionized water. The concentration of fluorescein was determined by electronic absorption using an extinction coefficient of 76,900 M^−1^cm^−1^ at 490 nm.^67^ The concentration of Atto Thio 12 was determined similarly, using an extinction coefficient of 1.1×10^5^ M^−1^cm^−1^ at 582 nm (Leica Microsystems). Small aliquots were prepared and stored at −20 °C. The frozen aliquots were thawed in a water bath at 4 °C on the day of the experiment.

### Vitamin C

Vitamin C solutions were prepared on the day of the experiment in distilled deionized water. Concentrations were assessed by electronic absorption using an extinction coefficient of 6,956 M^−1^cm^−1^ at 250 nm (pH-independent isosbestic point).^20^ The remaining solutions were discarded after the experiments.

### LC-photo-CIDNP molecules of interest

Stock solutions of Trp isotopologs (unlabeled Trp, Trp-U-^13^C,^15^N and QISP Trp), serotonin, melatonin, L-DOPA, epinephrine, indoxyl sulfate, Tyr, rizatriptan benzoate were prepared either in D_2_O or in the distilled deionized water and it was followed by filtration with 0.22 μm filter. Stock solutions of zolmitriptan, harmaline and reserpine were prepared in DMSO-d₆ due to their limited solubility in water. Concentrations were determined via electronic absorption spectroscopy using the following molar extinction coefficients: Trp isotopologs (5,540 M^−1^cm^−1^ at 280 nm)^68^, Tyr (1,280 M^−1^ cm^−1^ at 280 nm),^69^ L-DOPA (2,630 M^−1^cm^−1^ at 280 nm),^70^ serotonin (5,310 M⁻¹ cm⁻¹ at 280 nm),^68^ melatonin (6,300 M⁻¹ cm⁻¹ at 275 nm),^71^ epinephrine (2,754 M^−1^cm^−1^ at 280 nm),^72^ indoxyl sulfate (5037.5 M^−1^cm^−1^, measured in the lab), rizatriptan (89,683 M⁻¹ cm⁻¹ at 225 nm),^73^ and harmaline (18,527 M⁻¹ cm⁻¹ at 371 nm).^74^ In the case of zolmitriptan and reserpine, extinction coefficients are not available in the literature. Therefore, concentrations were estimated based on weighed mass. Small aliquots of each of the compounds were prepared and stored at −20 °C. The frozen aliquots in DMSO-d_6_ were thawed on a water bath at 35 °C and kept at room temperature during the experiments.

### Preparation of highly concentrated LC-photo-CIDNP dark (LED-off) samples

LC-photo-CIDNP experiments under dark conditions on highly concentrated samples were performed to estimate the enhancement factor (**ε**) (reported in **Fig. 2,3**). QISP Trp and the unlabeled Trp dark samples were prepared in a 10 mM potassium phosphate buffer (pH 7.2). At 600 MHz, for ^13^C RASPRINT and ^1^H PASS-W5ES dark experiments: 100 μM QISP Trp and 1 mM unlabeled Trp were used respectively. At 80 MHz, for ^13^C RASPRINT and ^1^H PASS-WET dark experiments: 1,430 μM QISP Trp and 1 mM unlabeled Trp were used respectively. 10 % v/v D_2_O was added to samples prepared for 600 MHz experiments. In the case of ^1^H PASS-WET or W5ES LC-photo-CIDNP samples, 500 µM DSS-d_6_ was added as an internal standard. A DSS-d_6_ in D_2_O spectrum was used as an external reference for all the ^13^C RASPRINT spectra.

### Preparation of LC-photo-CIDNP light (LED-on) samples in aqueous buffer

LC-photo-CIDNP experiments for ^13^C RASPRINT, ^1^H PASS-WET and ^1^H PASS-W5ES LC-photo-CIDNP under light conditions were prepared with the desired amounts of molecule of interest in either H_2_O- or D_2_O-based phosphate buffer (pH ∼7.2). Fluorescein was used for all the indole-scaffold-bearing compounds, whereas ATTO Thio 12 was used for phenolic scaffolds. Fluorescein and ATTO Thio 12 concentrations were optimized for different concentrations of molecules of interest (see Figure captions). In addition to the molecule of interest and the photosensitizer dye, 10 % v/v D_2_O (only at 600 MHz), 0.15 μM of GO, 0.1 μM of CAT enzymes and 2.5 mM D-glucose-d_12_ were added. VC was added only for low-μM samples (see Figure captions). 500 μM of DSS as an internal standard was added for the ^1^H PASS-WET and ^1^H PASS-W5ES LC-photo-CIDNP samples. The light experiments were carried out within less than 20 min after adding the D-glucose-d_12_. The power of the ^13^C-decoupler for the labeled ^13^C experiments was optimized using a 50 mM sample of uniformly ^13^C labeled glucose. The optimized decoupling power of GARP4 was 0.07384 W.

### Preparation of LC-photo-CIDNP samples in *E. coli* cell like medium

The S30 *E. coli* cell-like medium was prepared from BL21(DE3) cells transformed with a pET11d plasmid carrying the wild type drkN SH3 gene, following a modified version of a published procedure^75^ (also graphically illustrated in **Fig. 6** of the main manuscript). Cell cultures were grown overnight and harvested by centrifugation at 6,000 g for 15 min at 4 °C. The cell pellet was then resuspended in an ice-cold lysis buffer containing 50 mM Tris-HCl (pH ∼7.5), 2 mM EDTA, and a protease inhibitor (Pierce, A32965). The cells were then lysed via sonication with a Fisher Scientific FB505 device (500-Watt, 20 kHz) equipped with a tubular probe (Ultrasonic Convertor, model CL4) for 4 min (1 s-on/1 s-off and 65% vertical-amplitude motion) at 4 °C. The lysed cells were centrifuged at 30,000 g for 15 min at 4 °C to eliminate insoluble cell debris and fragmented genomic DNA. The resulting supernatant, containing the soluble protein fraction, was collected and used as the cell-like medium in LC-photo-CIDNP experiments. The 5-fold diluted cell-like medium (upon addition of 5-fold v/v 10 mM potassium phosphate) was mixed with the required amounts of QISP Trp or unlabeled Trp in 10 mM potassium phosphate buffer (pH 7.2). Fluorescein was used as photosensitizer dye (see **Fig. 6** caption for concentrations) and 0.15 μM of GO, 0.1 μM of CAT enzymes and 2.5 mM D-glucose-d_12_ were employed for oxygen-scavenging purposes.

### Preparation of LC-photo-CIDNP light samples in human serum

LC-photo-CIDNP samples were prepared upon mixing 5-fold diluted human serum, 10 µM epinephrine, 10 µM ATTO Thio 12, 0.15 μM GO, 0.1 μM CAT, 2.5 mM D-glucose-d₁₂, and 500 μM DSS-d₆ as internal standard. The human-serum stock solution was aliquoted into small fractions and stored at −20 °C. Freshly thawed aliquots were used for every experiment, and all remaining portions were discarded.

### 1D LC-photo-CIDNP NMR data collection at a high field (600 MHz)

All the high-field conventional NMR and LC-photo-CIDNP NMR experiments were carried out on an Avance III HD NMR spectrometer (Bruker Biospin Corp.) equipped with a 5 mm ^1^H[^19^F/^13^C/^15^N] 600 MHz triple-resonance cryogenic probe fitted with a z-gradient, and TopSpin 3.7.0 software. For LC-photo-CIDNP experiments, illumination was delivered through a 1.5 mm diameter, 4 m-long optical fiber (POF, Prizmatix, Holon, Israel). When fluorescein was used as the dye, a UHP-mic-LED-450 source (a.k.a., UHP-LED-blue; Prizmatix, Holon, Israel) with an emission maximum at 466 nm and 0.55 W power at the fiber tip was employed. When ATTO Thio 12 was used as the dye, a UHP-T-545-SR-LED source with emission maximum at 545 nm and 0.68 W power at the fiber tip was employed. The fiber tip was inserted into a 4 mm diameter NMR tube, which was then placed into a 5 mm diameter NMR tube which had the sample (**Fig. 1B**). ^13^C RASPRINT (light and dark) experiments in **Fig. 2** used 4,096 total data points, an acquisition time of 0.2048 s, a recycle delay of 0.5 s and an LED irradiation time of 1 s per scan. Note the ^1^H^α^ splitting due to the J-coupling to ^13^C^α^ was removed via GARP4 decoupling during acquisition. ^1^H PASS-W5ES LC-photo-CIDNP experiments in **Fig. 3** included 4,096 total data points, an acquisition time of 0.2138 s, a recycle delay of 1 s and an irradiation time of 1 s per scan. A much longer than usual recycle delay of 5 s was used for the dark experiments (unless stated otherwise) which were used to determine the enhancement factors (**ɛ**) both for ^13^C RASPRINT and ^1^H PASS-W5ES LC-photo-CIDNP. A long recycle delay of 5 s ensures complete spin-lattice relaxation between two successive scans.^19^ All spectra at 600 MHz were processed with MNova (version 15.1.0) upon zero-filling to 65,536 complex point and employing 5 Hz exponential decay as the window function. A DSS-d_6_ in D_2_O spectrum was used as external reference.

### 1D LC-photo-CIDNP NMR data collection at a low field (80 MHz)

The low-field conventional NMR work and LC-photo-CIDNP experiments were performed on a Fourier-80 Benchtop NMR equipped with [^1^H/^13^C/^19^F] room-temperature probe, external lock (no deuterated solvent was required) and TopSpin 4.4.0 software. For LC-photo-CIDNP experiments, illumination was delivered through a 1.5 mm diameter, 2 m long optical fiber (POF, Prizmatix, Holon, Israel). When fluorescein was used as the dye, a UHP-mic-LED-450 source (emission maximum at 453 nm and 0.55 W power at the fiber tip) was employed. When ATTO Thio 12 was used as the dye, a UHP-mic-545 LED source (emission maximum at 545 nm and 0.68 W power at the fiber tip) was used. The fiber tip was inserted into a 4 mm diameter NMR tube, which was then placed into a 5 mm diameter NMR tube which had the sample **(Fig. 1B)**. For ^13^C RASPRINT light experiments in **Fig. 2** and **Fig. 6**, 784 total data points, an acquisition time of 0.2979 s, a recycle delay of 0.5 s and an LED irradiation time of 1 s per scan were used, unless otherwise stated. Note the ^1^H^α^ splitting due to the J-coupling to ^13^C^α^ was removed using GARP4 decoupler during acquisition. ^1^H PASS-WET LC-photo-CIDNP (light and dark) experiments used 4,096 total data points and an acquisition time of 1.7203 s, a recycle delay of 1 s and an LED irradiation time 1 s per scan were used. For ^1^H PASSWORD LC-photo-CIDNP experiments in **Fig. 6G**, 126 T_2_-filter loops (n=126, 79 ms filtering time), 1 s recycle delay, 4,096 total data points and 1.72 s acquisition time and 1 s LED irradiation time per scan were used. A much longer than usual recycle delay of 5 s was used for the dark experiments (unless stated otherwise) which were used to determine the enhancement factors (**ɛ**) for the same reason mentioned above. All spectra at 80 MHz were processed with MNova (version 15.1.0) upon zero-filling to 65,536 complex point and 1 Hz exponential decay and as the window function. A DSS-d_6_ in D_2_O spectrum was used as external reference.

### 2D LC-photo-CIDNP NMR data collection at a low field (80 MHz)

In 2D ^1^H-^1^H COSY LC-photo-CIDNP experiments (**Fig. 4F**), data were acquired on a Fourier-80 benchtop NMR spectrometer using the identical optical setup described above. Experiments were performed in QF 2D acquisition mode. Both light and dark LC-photo-CIDNP experiments used a sweep width of 3.0 ppm in both dimensions, with the carrier frequency (O1p) set to 7.360 ppm. A total of 1,024 data points were acquired in the direct dimension, while 128 total transients were collected in the indirect dimension, with 4 scans per transient. Spectra were processed in magnitude mode and zero-filled four-fold in the direct dimension and six-fold in the indirect dimension. A slightly shifted Gaussian apodization function was applied in both dimensions. The 2D data were processed with NMRPipe (v. 9.0.0-b108) and visualized with the NMRDraw and NMRView J (v. 2009.015.15.35) software packages.^76,77^

### 2D NMR data collection at high concentration at a low field (80 MHz)

2D ^1^H-^1^H COSY data were collected on a Fourier-80 benchtop NMR spectrometer on 1.4 mM serotonin sample (**Supplementary Fig. 9**). The experiments used a 3.0 ppm sweep width in both dimensions, with carrier frequency at 7.360 ppm for both. 1,024 total data points were used in the direct dimension while 256 transients were collected in the indirect dimension. 4 scans were collected per transient. The spectra were zero-filled 4-fold and 5-fold in the direct and indirect dimension, respectively. A slightly shifted Gaussian apodization function was applied to both dimensions. The 2D data were processed with NMRPipe (v. 9.0.0-b108) and visualized with the NMRDraw and NMRView J (v. 2009.015.15.35) software packages. ^77,76^

### SHARPER-enhanced ^1^H LC-photo-CIDNP data acquisition

The ^1^H *sel*-SHARPER pulse sequence^56^ was adapted for hyperpolarized data collection upon addition of LED on/off trigger commands preceding the radiofrequency pulses (**Fig. 5D**, top region). The interval between two consecutive 180° pulses (during acquisition) was optimized to 5 ms, for the serotonin aromatic resonances. The bandwidth of the selective 180° refocusing pulse was set to 84 Hz, with O1p at 7.0 ppm. Selective refocusing was achieved via the *gaus180r* shaped pulse (10.7 ms, 1.6 W). An SMSQ10.100 gradient pulse (G1) with a duration of 1 ms, 13% CTP, and a 400 μs gradient recovery delay was used. The imaginary component of the FID was removed using an AU program in TopSpin 4.4.0, as previously described.^55^ A total of 4,096 data points were used with 1.27 s acquisition time, 1 s recycle delay and 1 s LED irradiation time per scan. Processing parameters were identical to those employed for ^1^H PASS-WET LC-photo-CIDNP.

### Computational assessment of field-dependent geminate polarization of Trp isotopologs

We carried out applied magnetic field (B_0_) dependent calculations for both ^13^C^α^ and ^1^H geminate polarization of Trp-U-^13^C,^15^N, QISP Trp, and unlabeled Trp respectively (**Fig. 2C** and **Fig. 3B**). The computations were based on Adrian’s theoretical modelling of the radical pair mechanism. Equations and methods (see **Supplementary Information** for details) were as described.^18,19^ Hyperfine parameters used in these models are provided in **Supplementary Table 2**. All computations were performed using a custom Python 3.12 workflow running on the University of Wisconsin–Madison Center for High-Throughput Computing (CHTC).

