## Supplementary Information for "Hypersensitive Detection of Neurotransmitters in Biological Media by Optically Enhanced Benchtop NMR Spectroscopy"

[1] Equal contributors

#### Supplementary Text

**Tutorial on the basic mechanistic aspects of steady-state photo-CIDNP (and LC-photo-CIDNP) in liquids.** As pictorially outlined in the text of the main article (see also **Fig. 1C**), steady-state photo-CIDNP in solution proceeds via several key steps. The corresponding mechanism[1,2] is summarized below. First, the sensitizer dye is photoexcited and undergoes a transition from its ground singlet state $S_0$ to its photo-excited triplet state $S_1$. In most cases, the latter $S_1$ state is not sufficiently long-lived to collide with molecules of interest, in a dilute solution of the latter (e.g., ca. 10 μM or less). Therefore, the dye undergoes intersystem crossing (ISC) to generate photoexcited triplet states ($T_0$, $T_+$, $T_-$) within 0.1 ns – 1 μs. Collision with the molecule of interest is rapidly followed by electron transfer (overall timescale: 10 ps – 1 μs): note that the collision is a bimolecular process that depends on the concentration of the molecule of interest. Due to the electron transfer process, the dye is typically reduced, and the molecule of interest is correspondingly oxidized, generating triplet radical pairs ($T_0$, $T_+$, $T_-$). The energy levels of the triplet and $T_+$ and $T_-$ states of the radical pair are highly separated relative to the corresponding singlet state energy level. Therefore, the $T_+$ and $T_-$ radical pairs eventually dissociate.

On the other hand, the nearly degenerate $T_0$ and singlet (S) states of the radical pair (denoted as $D^{\cdot-}M^{\cdot+}$ in **Fig. 1C**) are allowed to undergo coherent triplet-singlet (TS) mixing, in the presence of a magnetic field of significant intensity (e.g., > 0.5 T) and when the elements of the radical pair are at a sufficiently large distance (usually a few Å) for the exchange interaction between the two unpaired electrons to be negligible. Under these conditions, the radical pair undergoes TS mixing.[3] The frequency of this TS mixing ($\omega_{TS}$) of the $T_0$ radical pair is nuclear-spin-dependent and plays a critical role in determining the non-Boltzmann spin distribution in the final hyperpolarized products of a steady-state photo-CIDNP experiment, as further discussed below. Generation of the singlet (S) state of the $T_0$ radical pair is followed by immediate back electron transfer from the dye to the molecule of interest. This process (i.e., regeneration of the diamagnetic molecule of interest while generating a non-Boltzmann spin distribution) is known as geminate recombination. In contrast, $T_0$ triplet-state radical pairs experiencing slow TS-mixing rates undergo "escape" and dissociate into their individual components before any significant extent of triplet-singlet mixing takes place.

The molecules of interest originating from the "escaped" $T_+$, $T_-$ and $T_0$ radical pairs are typically sufficiently long-lived, in dilute solution, to undergo paramagnetic relaxation during which the non-Boltzmann distribution of nuclear spin states is lost. Some of the paramagnetically relaxed escaped molecules, upon random collision, statistically generate some additional $T_0$ radical pairs, which ultimately give rise to some more polarization via a process similar to the one described in the previous paragraphs. Ultimately, the observed hyperpolarization is the net result of the above pathways.

Importantly, as already mentioned, the frequency of TS-mixing of the $T_0$ radical pair ($\omega_{TS}$) depends on state of the nuclear spins interacting with the unpaired electrons (via rotationally averaged isotropic hyperfine coupling constants). The $\omega_{TS}$ of either of the nuclear spin states (α or β), of a hypothetical molecule composed of a single spin ½ nucleus, including all other nuclear-spin configurations (χ) of the radical pair, is

$$\omega_{TS,\ \alpha\ \mathrm{or}\ \beta,\ \chi} = \frac{1}{2}\left[\left(g_D - g_M\right)\beta_e B_0 \hbar^{-1} + \sum_{i=0}^{a} m_i A_i - \sum_{j=0}^{b} m_j A_j\right]\ , \qquad \text{(S1)}$$

where, $g_D$ and $g_M$ are the electronic g-factors of the dye (D) and the molecule of interest (M), respectively, $\beta_e$ is Bohr magneton of electron (in J $T^{-1}$), $\hbar$ is reduced Planck's constant (in J.s), $m_i$ is the magnetic spin quantum number, A is the hyperfine coupling constant, i and j are the nuclear spin states of D and M, respectively, with the chosen nuclear spin configuration, a is the number of NMR-active nuclei of D, b is the number of NMR-active nuclei of M minus 1, where the "minus 1" arises from the fact that the nucleus of interest is excluded. The A values, which are typically reported in milli Tesla (mT), need to be converted to Hz, upon taking the electron gyromagnetic ratio ($\gamma_e$= 28.02 GHz/T) into account.[4]

The population difference between any two nuclear configurations α, χ and β, χ in a freely diffusing radical pair can be expressed as

$$p_{\alpha,\chi} - p_{\beta,\chi} = p\sqrt{\tau_D}\left(\sqrt{|\omega_{TS,\alpha,\chi}|} - \sqrt{|\omega_{TS,\beta,\chi}|}\right) \quad , \qquad \text{(S2)}$$

where $p_{\alpha \text{ or } \beta,\chi}$ are the population in α, χ and β, χ nuclear-spin configurations, $\tau_D$ is the lifetime of the radical pair. The latter is the average time the two components of the radical pair spend close to each other within a "solvent cage", before they diffuse apart, and p is the normalization factor to ensure that the sum of all possible configurations equals 1. The geminate polarization of the nucleus of interest in Trp ($P_0$) therefore equals the difference between the sum of all populations with configurations where the nucleus of interest is in the α spin-state and the sum of all populations with configurations where the nucleus of interest is in the β spin-state. Therefore, the net polarization ($P_0$) of the geminate recombination product is

$$P_0 = \sum p_{\alpha,\chi} - \sum p_{\beta,\chi} = p\sqrt{\tau_D}\left(\sum\sqrt{|\omega_{TS,\alpha,\chi}|} - \sum\sqrt{|\omega_{TS,\beta,\chi}|}\right) \quad . \qquad \text{(S3)}$$

The normalization factor p is defined as

$$1 = \sum p_{\alpha,\chi} + \sum p_{\beta,\chi} = p\sqrt{\tau_D}\left(\sum\sqrt{|\omega_{TS,\alpha,\chi}|} + \sum\sqrt{|\omega_{TS,\beta,\chi}|}\right) \quad , \qquad \text{(S4)}$$

In summary, $\tau_D$ contributes to the extent of observed polarization so that, for instance, longer $\tau_D$*s* lead to higher polarization, within the geminate recombination product. Towards this end, photosensitizer dyes like fluorescein, which are characterized by a long photoexcited triplet-state lifetime (~20 ms in the absence of oxygen at low concentration), are ideal. A long triplet-state lifetime increases the likelihood of productive collisions with the molecule of interest, while the dye is still in its photoexcited triplet state, thereby promoting formation of triplet-state radical pairs. In addition, dyes bearing a long-lived triplet state in solution are more likely to collide with the molecule of interest while still in the triplet state, contributing to the generation of geminate polarization. The $\tau_D$ parameter can be determined according to

$$\tau_D = \frac{(R_D+R_M)^2}{D_D+D_M} \quad , \tag{S5}$$

where $R_D$ and $R_M$ are the van der Waals radii of the radicals of the dye and the molecule of interest, respectively. $D_D$ and $D_M$ are the translational diffusion coefficient of the dye and the molecule of interest, respectively.

As an example, the TS mixing frequency of the "0th" nucleus of interest in the α and β states, given that χ represents the configurations of all the other nuclei in the dye and molecule of interest, can be expressed as

$$\omega_{TS,\alpha,\ \chi} = \frac{1}{2}\ \{(g_D\text{-}g_M)\beta_e B\hbar^{-1}+\sum_{i=0}^{a} m_i\, A_i - \sum_{j=0}^{b} m_j\, A_j - \frac{1}{2}\ A_0 \quad , \tag{S6}$$

$$\omega_{TS,\beta,\ \chi} = \frac{1}{2}\ \{(g_D\text{-}g_M)\beta_e B\hbar^{-1}+\sum_{i=0}^{a} m_i\, A_i - \sum_{j=0}^{b} m_j\, A_j + \frac{1}{2}\ A_0 \quad . \tag{S7}$$

The $P_0$ simulations were carried out with a custom-built python script (Python v 3.8) using equations (S3) and (S4). All calculations were performed at UW-Madison Center for High-Throughput Computing (CHTC). The following parameters were used for the calculations : g(Trp·+) = 2.0027,{Kiryutin, 2007 #16} g(Fl·-) = 2.003077,[5] $D_D$ = 4.2 × 10-6 $cm^2s^{-1}$, $D_M$ = 6.592 × 10-6 $cm^2s^{-1}$, $R_D$ = 4.4 Å and $R_M$ = 4.2 Å. The hyperfine coupling constants of Trp and Fl were derived from the literature.[5,6] Since the hyperfine coupling constant of a nucleus is proportional to its gyromagnetic ratio, the hyperfine coupling constants of specific deuterons ($^2H$) in QISP Trp were obtained by scaling the corresponding $^1$H hyperfine coupling constants by the ratio $\frac{\gamma_{1_H}}{\gamma_{2_H}}$, as described.[7] Additional details are provided in **Supplementary Table 2**. Perhaps needless to write, the hyperfine coupling constant of $^{12}$C nuclei was regarded as 0, and the natural-abundance contribution of $^{13}$C nuclei (~1%) in QISP Trp was neglected. The hyperfine coupling constants of the Trp primary-amine nitrogen was not included in the calculations, given that its value is not available in the literature, to the best of our knowledge. This omission is expected to introduce no appreciable simulation misestimates, as shown in previous control calculations.[5] In this work, we simulated the geminate polarization of all natural-abundance protons of Trp as a function of applied magnetic field (**Fig. 3A**).

**Simulation of NMR spectral properties and spectra in the absence of a cryogenic probe.** Experimentally acquired cryoprobe spectra ($^{13}$C RASPRINT or $^1$H PASS-W5ES) at 600 MHz were used as the starting datasets. The average noise level ($n$) was estimated from the standard deviation ($\sigma$) of a resonance-free baseline region (8–10 ppm). To approximate the full peak-to-peak noise range, $n \pm 3\sigma$ interval (corresponding to ~99.7% of the Gaussian noise distribution) was considered, so the noise was effectively scaled by a factor of $6\sigma$. To simulate the performance of a room-temperature probe, $6\sigma$ was further scaled by a factor of 3.5, corresponding to the typical 3–4-fold noise reduction provided by cryogenic probes.[8] Gaussian noise bearing the above-described scaled amplitude was added to the original dark and light spectra acquired in the presence of a cryogenic probe. The resulting spectra correspond to simulated non-cryoprobe data sets. All computations and noise simulations were performed with a custom-made Python script (see **Appendix**) generated on a MacOS system.

**Optimization of LED irradiation times in benchtop-NMR LC-photo-CIDNP experiments.** In order to identify an optimal irradiation time for both $^{13}$C RASPRINT and $^1$H PASS-WET LC-photo-CIDNP experiments performed on QISP and unlabeled tryptophan, respectively, several LED durations were experimentally probed. Namely, the LED durations tested for $^{13}C$ RASPRINT with QISP Trp were 100 ms, 150 ms, 200 ms, 250 ms, 300 ms, 400 ms, 500 ms, 600 ms, 700 ms, 800 ms, 900 ms, 1 s, 2 s, 3 s, and 4 s. To limit sample degradation due to the continuous LED irradiation, two fresh samples were prepared for each data point, the first sample had 100 ms through 2 s LED durations and the second sample had 2 s, 3 s and 4 s. Each duration that was tested had 4 scans and three independent repeats, with six total samples used. The optimal LED irradiation duration was found to be 1 s (**Supplementary Fig. 1A**). The LED durations tested for $^{1}H$ PASS-WET LC-photo-CIDNP with unlabeled Trp were 200 ms, 250 ms, 300 ms, 350 ms, 400 ms, 500 ms, 600 ms, 700 ms, 800 ms, 900 ms, 1 s, 2 s, 3 s, 4 s, 5 s, and 6 s. Same as with $^{13}C$ RASPRINT, two samples were used for each data point, the first sample has the LED durations 200 ms through 2 s, and the second sample had the LED durations 3 s, 4 s, 5 s, and 6 s. Each LED duration tested had 4 scans and 3 total replicates, totaling six samples. The optimal LED irradiation duration was found to be 1 s upon fitting the data points to the equation (S8) (**Supplementary Fig. 1B**). Each sample contained 100 μM of the respective tryptophan isotopolog, 25 μM fluorescein, 10 mM potassium phosphate buffer (pH ~ 7.2), 0.15 μM of GO, 0.1 μM of CAT, 2.5 mM D-glucose-$h_{12}$ (for $^{13}C$ RASPRINT) or 2.5 mM glucose-$d_{12}$ (for $^{1}H$ PASS-WET). All samples were prepared in water-based buffer. To limit variation in samples, the same NMR tube was used for all the experiments, and all independent experiments for each pulse sequence and isotopolog were conducted on the same day.

The intensity of steady-state photo-CIDNP hyperpolarization, for transitions between nuclear spin states m and n ($I_{mn}$), can be expressed as[9]

$$I_{mn} = I^0_{mn} + CrT_1(P_m - P_n)\left(1 - \exp\left(-\frac{\tau}{T_1}\right)\right) \ , \qquad (S8)$$

where $I^0_{mn}$ is the NMR signal intensity for m-to-n transition in the absence of photo-CIDNP (no LED irradiation), C is the proportionality constant, r is the rate of formation of radical pairs (assumed to be time independent), $T_1$ is the longitudinal relaxation time of the diamagnetic nucleus of interest, and $P_m$, $P_n$ are the total nuclear spin-dependent recombination probabilities of the radical pair and τ is the duration of the LED irradiation. For optimal sensitivity, τ should be in the order of $T_1$. A single-exponential function (equation S8) was used to fit the data points. The experimental $T_1$ values (**Supplementary Table 7**) were used in the fitting, with two fitting parameters: $I^0_{mn}$ and $Cr\ (Pm - Pn)$. For the aliphatic region, the average $T_1$ of $^1H^{\beta1}$ and $^1H^{\beta2}$ (0.52 s) was used. For the aromatic region, the average $T_1$ of $^1H^{\delta1}$ and $^1H^{\eta2}$ (1.87 s) was used and the $T_1$ value for $^1H^{\varepsilon3}$ (1.79 s) and $^{13}C^{\alpha}$ (1.37 s) were used as measured in $H_2O$-based phosphate buffer.

**$T_1$ measurements on $^1H$ and $^{13}C^{\alpha}$ nuclei by benchtop NMR spectroscopy**. For $^1H$ $T_1$ measurements at 600 MHz, 5 mM unlabeled Trp in 10 mM $H_2O$-or-$D_2O$-based potassium phosphate buffer (pH ~7.2) and 500 μM DSS was used. An inversion recovery sequence coupled to WET as solvent suppression at 80 MHz was used **(Supplementary Fig. 10).** The $^1H$ $T_1$ inversion recovery experiments at 80 MHz both in $D_2O$ and $H_2O$, used a recycle delay of 20 s, an acquisition time of 1.72 s and 4,096 total points. Inversion recovery delays for the experiments in $D_2O$ were set to 10 ms, 50 ms, 100 ms, 250 ms, 500 ms, 800 ms, 1.5 s, 3 s, 5 s and 10 s, with 8 scans acquired per delay. Inversion recovery delays for the experiments in $H_2O$ were set to 1 ms, 5 ms, 60 ms, 100 ms, 200 ms, 500 ms, 1 s, 2 s, 4 s, 8 s, 10 s, with 64 scans acquired per delay (see discussion on the pulse sequence later). A modified $^{13}C^{\alpha}$ RASPRINT like pulse sequence as described in Li *et al*. was used to measure $^{13}C^{\alpha}$ $T_1$.[10]

A 1.34 mM QISP Trp in 10 mM $H_2O$-based potassium phosphate buffer (pH ~7.2) was used for $^{13}C^{\alpha}$ $T_1$ measurements. A recycle delay of 10 s, an acquisition time of 0.288 s and 784 total data points were used. Inversion recovery delays were set to 1 ms, 100 ms, 300 ms, 800 ms, 1 s, 1.5 s, 1.8 s, 3 s, 5 s, 7 s and 10 s. 1600 scans were collected for each delay. All $T_1$ measurements were repeated twice at both fields. The data were fit according to

$$I(t)=I(\infty)(1-2e^{-\tau/T_1}) \quad , \qquad \text{(S9)}$$

where I (t) is the area of the resonance of interest at a particular inversion recovery delay time τ, I (∞) is the area of the resonance of interest at equilibrium magnetization, $T_1$ is the longitudinal relaxation time. $^1H$ $T_1$ in $D_2O$ data fitting was done in TopSpin 4.5.0 using Dynamics $T_1/T_2$ module and $^1H$ and $^{13}C^{\alpha}$ $T_1$ fitting in $H_2O$ at 80 MHz were performed in Kaleidagraph. The area with the longest inversion-recovery delay was taken as I (∞) in the fits (assuming the magnetization to have fully relaxed to equilibrium), and $T_1$ was used as the only adjustable parameter.

**Chemical shift validation of LC-photo-CIDNP resonances of amino acids, neurotransmitters, biomarkers and pharmaceuticals.** Chemical shifts of L-DOPA, tyrosine, epinephrine, melatonin, serotonin, indoxyl sulfate, harmaline and reserpine were assigned according to literature (also see **Supplementary Fig. 5,6).**[11,12,13,14,15-18]

Rizatriptan benzoate chemical shifts were not available in the literature and therefore assigned using zolmitriptan as a reference, to the best of our abilities. We also note that the strong-coupling regime inherent to low-field NMR occasionally limited our ability to resolve individual coupling constants. In the $^1H$ NMR spectrum of zolmitriptan shown in the **Supplementary Fig. 5D**, $^1H^{\eta 2}$ appeared as doublet of doublet (7.13 ppm, J=8.8 Hz for ortho coupling with $^1H^{\eta 1}$ and 1.6 Hz for meta coupling with $^1H^{\varepsilon 2}$), $^1H^{\delta 1}$ appeared as a singlet (7.31 ppm), $^1H^{\eta 1}$ appeared as a doublet (7.45 ppm, J=8.24 Hz for ortho coupling with $^1H^{\eta 2}$) and $^1H^{\varepsilon 2}$ appeared as a singlet (7.52 ppm). See

Neelakandan *et al.* for the complete aromatic assignments. Note that the resonance at 7.52 ppm integrates to approximately twice the area of the 7.45 ppm resonance, indicating that the downfield component of the $^1H^{\eta 1}$ doublet overlaps with the $^1H^{\varepsilon 2}$ (7.52 ppm) resonance. We also noticed that under LC-photo-CIDNP conditions, rizatriptan and zolmitriptan exhibit nearly identical splitting patterns and scalar couplings. So, with the knowledge of zolmitriptan assignments, we assigned the rizatriptan aromatic resonances. In addition, rizatriptan used in this work contained a benzoate counterion, which introduced extra aromatic resonances in the $^1H$ NMR spectrum (**Supplementary Fig. 5B**). So, a benzoic acid reference spectrum at pH ~7.2 was thus collected (**Supplementary Fig. 5B, reference spectrum)**, which helped in excluding the benzoic acid resonances within 7.78–7.95 ppm and 7.49–7.54 ppm ranges. Rizatriptan $^1H$ NMR aromatic chemical shift assignments: $^1H^{\eta 2}$ appeared as doublet of doublet (7.21 ppm, J=7.2 Hz for ortho coupling with $^1H^{\eta 1}$ and 1.6 Hz for meta coupling with $^1H^{\varepsilon 2}$), $^1H^{\delta 1}$ appeared as a singlet (7.36 ppm), $^1H^{\eta 1}$ appeared as a doublet (7.54 ppm, J = 8.0 Hz for ortho coupling with $^1H^{\eta 2}$), $^1H^{\varepsilon 2}$ appeared as a singlet (7.65 ppm), $^1H^{\lambda 1}$ appeared as a singlet (8.04 ppm) and $^1H^{\kappa 2}$ appeared as a singlet (8.58 ppm).

**Potential role of DMSO in enhancing LC-photo-CIDNP signal-to-noise ratios (S/N) of highly nonpolar compounds.** A full mechanistic analysis of the effect of DMSO on LC-photo-CIDNP intensities is beyond the scope of this work, but our observations point to at least three plausible contributing mechanisms – (i) Reserpine and harmaline are the only two compounds for which the addition of DMSO increased the LC-photo-CIDNP signal-to-noise ratio. Both molecules are highly hydrophobic (logP for reserpine = 3.32 and logP for Harmaline = 1.67) and are only sparingly soluble in aqueous buffer; the introduction of DMSO likely prevents the formation of any aggregates and improves molecular dispersion. This would likely enhance the efficiency of dye–molecule encounters and thereby promote productive radical-pair formation. (ii) DMSO has a viscosity roughly 2.3-fold higher than water. Increased viscosity slows the translational diffusion down and according to radical-pair theory it can modify geminate re-encounter probabilities and was proven to increase the LC-photo-CIDNP polarization.[19] (iii) DMSO is also known to influence the redox environment in solution, which could potentially alter the electron-transfer properties of both the molecule and the photosensitizer dye,[20] potentially affecting the efficiency of radical-pair formation, therefore the resulting CIDNP signal.

**Product-operator description of COSY LC-photo-CIDNP.** The COSY LC-photo-CIDNP spectrum shown in **Fig. 4F** of the main manuscript is not symmetric, in contrast to a conventional COSY NMR spectrum. This asymmetry can be rationalized by examining the product-operator evolution underlying the LC-photo-CIDNP experiment. Consider two scalar-coupled protons, $H_1$ and $H_2$, where only one proton ($H_2$) is directly hyperpolarized *via* LC-photo-CIDNP. The evolution of the initial magnetization terms $H_{1z}$ and $H_{2z}$ throughout the COSY LC-photo-CIDNP pulse sequence during actual scans **(Supplementary Fig. 8)** is shown below:

$$\boldsymbol{H_{1z} + H_{2z} \xrightarrow{LED\ (t_L)} H_{1z} + \varepsilon H_{2z} \xrightarrow{(\pi/2)_x - t_1 - (\pi/2)_x}}$$

$$\boldsymbol{(\varepsilon - \varepsilon')H_{2y}\sin(\Omega_2 t_1) + H_{1y}}\sin(\Omega_1 t_1) + \boldsymbol{\varepsilon' 2H_{2z}H_{1x}sin(\Omega_2 t_1)} + \ \boldsymbol{2H_{2x}H_{1z}}\sin(\Omega_1 t_1) + \text{non}$$

-observables.

Here, $\varepsilon$ denotes the LC-photo-CIDNP enhancement factor for $H_2$ prior to the first RF pulse, $\varepsilon'$ is the LC-photo-CIDNP enhancement factor for $H_2$ transferred from $H_1$ at the start of the acquisition, $t_1$ is the evolution time in the indirect dimension, and $\Omega_m$ represents the chemical shift evolution of the $m^{th}$ proton. At the start of acquisition, two of the four observable terms ($\varepsilon H_{2y}\sin(\Omega_2 t_1)$ and

$\varepsilon 2H_{2z}H_{1x}\sin(\Omega_2 t_1)$) are enhanced by LC-photo-CIDNP. At the low concentrations used in this study, the non-hyperpolarized terms i.e., $H_{1y}\sin(\Omega_1 t_1)$ and $2H_{2x}H_{1z}\sin(\Omega_1 t_1)$, are effectively undetectable. Consequently, during the evolution period ($t_1$), only directly hyperpolarized protons ($H_2$) contribute to the signal, and the projection onto the indirect dimension resembles a conventional 1D $^1$H PASS-WET LC-photo-CIDNP spectrum. In contrast, in the direct dimension, both directly polarized ($H_2$) and cross-polarized ($H_1$) protons are observed.[21]

**Basic features of $^1$H WET-like inversion recovery experiment for $^1$H $T_1$ measurements.** We used $^1$H WET-like inversion recovery pulse sequence to measure $^1$H $T_1$ values of unlabeled Trp in water-based buffer (pH ~7.2). Here, note that $^1$H $T_1$ values in $H_2O$ were used for fitting the data in **Supplementary Fig. 1A.** The $^1$H $T_1$ values of Trp in $D_2O$ are different from the ones in $H_2O$, since $D_2O$ is ~20-30% more viscous than $H_2O$ and deuterium has a much smaller gyromagnetic ratio than proton, henceforth $D_2O$ as a solvent can – (i) reduce the tumbling of molecule of interest, thus affecting the rotational time and (ii) reduce the intermolecular interaction between the proton of Trp and proton of water and each one of these can potentially change the $T_1$ values.[22,23] So, to measure the $T_1$ values in $H_2O$, the standard WET sequence was modified by introducing a non-selective 180° inversion pulse at the beginning of the experiment to invert the longitudinal magnetization **(Supplementary Fig. 10)**. This was followed by the inversion recovery delay and semi-selective WET pulses used for solvent suppression and, subsequently, signal acquisition. Henceforth, the effective magnetization recovery period consisted of the user-defined inversion-recovery delay plus the duration of the WET module (i.e., 100 ms, excluding the detection pulses).

**Basic features of $^1$H PASSWORD LC-photo-CIDNP pulse sequence**. $^1$H PASSWORD is an acronym for (**P**ulse-**A**cquire with **S**olvent **S**uppression via **W**ET **O**vercoming **R**elaxation-related **D**efects). $^1$H PASSWORD sequence shown in the **Fig. 6F** of the main manuscript uses the following delays: $\tau_1$ is 2 ms, $\tau_{WET}$ is (10 ms - $\tau_1$), $\tau_2$ is 292 ms, $\tau_3$ is 300 ms and $\tau_4$ is 290 ms. n is the number of loops used. The 4 bell-shaped curves denote 90-degree shaped pulses (sinc1.1000), each one of the shaped pulses is followed by a gradient pulse (SMSQ10.100) at 80%, 40%, 20%, 10% respectively. The duration level for each gradient was set to 1 ms. The phase cycling for $\phi_{rec}$ is x, x, -x, -x, y, y, -y, -y; $\phi_1$= x; $\phi_2$= y; $\phi_3$= x, x, -x, -x, y, y, -y, -y; $\phi_4$= y, -y, y, -y, x, -x, x, -x.

**Basic features of $^1$H PASS-CPMG-T$_2$-filter LC-photo-CIDNP pulse sequence**. This sequence was used for the reference experiments shown in **Fig. 6** of the main manuscript. To ensure stable performance, a set of dummy scans ($n_d$) was executed at the beginning; during these dummy scans, the LED was gated off to prevent unnecessary dye or sample degradation.[24] The dummy scans were followed by an identical set of pulses, the actual scans (ns), in which the LED was gated on in the beginning. LED control was automated through a TTL command in the pulse program **(Supplementary Fig. 13)**.[24] Adding a small number of dummy scans also eliminated analog-to-digital converter (ADC) overflow errors which are often encountered during data acquisition. $n_d$ can be set to any chosen number of iterations, and for all experiments reported here, $n_d$ was set to 4. The phase cycle used for this sequence: $\phi_1$ = x, x, -x, -x, y, y, -y, -y; $\phi_2$ = y, -y, y, -y, x, -x, x, -x; $\phi_{rec}$ = x, x, -x, -x, y, y, -y, -y.

### Supplementary Tables

**Supplementary Table 1.** Comparison between results achieved with different hyperpolarization strategies adapted to date in low-field NMR spectroscopy performed on benchtop NMR spectrometers.

| | Parameter | SABRE | dDNP | LC-photo-CIDNP |
|---|---|---|---|---|
| **General features of the technique** | **Molecular requirements** | Molecules able to reversibly bind metal catalyst | None | Aromatic compounds, reducing (low$E^0$)[a] molecules |
| | **Effective polarization time** | < 4 hrs ($x_p$ = 98% at < 25 K)[b, c]<br>< 1 hrs ($x_p$ = 50% at 77 K)[b, d] | - | - |
| **Specific studies (see references below for details)** | **Suitable molecule of interest** | N-containing heterocycles, amines | $^{13}$C-labeled metabolites and ligands | Trp and Trp derivatives, heteroaromatic compounds |
| | **Solvent** | MeOH, $CD_3OD$, EtOH | DMSO, $H_2O$, $D_2O$ | aqueous buffer, $D_2O$, urine |
| | **Polarization temperature (K)** | 298 K | 1.2: Ref. 29<br>77 K: Ref. 30 | 298 K |
| | **NMR-measurement temperature (K)** | 298 K | 77 K and 298 K | 298 K |
| | **Sample concentration**[e] | 40 µM–260 mM | 30 µM: Ref. 32<br>3 M: Ref. 33 | 100 nM–40.5 µM |
| | **Polarizing agent or photosensitive dye** | Ir catalyst<br>(0.1 mM – 5.2 mM) | TEMPOL<br>25 mM: Ref. 32<br>50 mM: Ref. 33 | Photosensitive dye<br>(2.5 µM-270 µM) |
| | **Detected nucleus** | $^{1}$H, $^{13}$C, $^{15}$N, $^{19}$F | $^{13}$C and $^{1}$H | $^{1}$H and $^{19}$F |
| | **$B_0$/$^1$H freq. (T/MHz)** | 1–1.4 T / 43-60 MHz | ~1.88 T / 80 MHz | 1.88 T / 80 MHz |
| | **Polarization buildup time** | 10 s – 1.5 min | 1 s ~ 10 min[f] | 0.2 s ~ 6 s |
| | **Enhancement factor (ε)** | ~ 360 ($^{19}$F)<br>~ 17,000 ($^{1}$H)<br>~ 45,000 ($^{13}$C)<br>~ 320,000 ($^{15}$N) | ~ 100 ($^{1}$H) | ~ 313 ($^{13}$C)<br>~ 465 ($^{19}$F)<br>1300 ($^{1}$H) |
| | **Percent polarization ($P_{\%}$)** | ~ 0.12% ($^{19}$F)<br>~ 4% ($^{13}$C)<br>~ 5.9 % ($^{1}$H)<br>~ 12% ($^{15}$N) | ~ 0.13% ($^{1}$H): Ref. 33 | ~ 0.09% ($^{19}$F)<br>~ 0.202% ($^{1}$H)<br>~ 0.837% ($^{13}$C) |
| | **Need for sample freezing** | NO | YES | NO |
| | **Need for sample transfer to a different device** | YES: All others<br>NO: Ref. 27[g] | YES: Ref. 32<br>NO: Ref. 33 | NO |
| | **References** | [25-32] | [33,34] | [35-38] |

[a] $E^0$ denotes the standard-state redox potential.
[b] $x_p$ is the para-$H_2$ mole fraction.
[c] < 4 hrs: time required to reach 98% para-$H_2$ polarization at temperatures below 25 K ([39]).
[d] < 1 hrs: time required to reach 50% para-$H_2$ polarization at 77 K ([40]).
[e] This is the lowest concentration tested in the publication.
[f] The reported 10 min per cross-polarization (CP) cycle includes both the CP step and the subsequent $^1$H relaxation period (33).
[g] *In situ* hyperpolarization by continuous para-$H_2$ bubbling directly into the NMR tube; no sample transfer required.

**Supplementary Table 2**. Hyperfine coupling constants (A) used in geminate-polarization simulations.

| Nucleus | A value (mT) | Ref. |
|---|---|---|
| **Trp•+ $^{13}C^{\alpha}$** | 0.5643[a] | [6] |
| **Trp•+ $^{13}C^{\beta}$** | -0.512 | [6] |
| **Trp•+ $^{13}C'$** | -0.045 | [6] |
| **Trp•+ $^{13}C^{\delta1}$** | -0.227 | [6] |
| **Trp•+ $^{13}C^{\gamma}$** | 1.254 | [6] |
| **Trp•+ $^{13}C^{\delta2}$** | -0.891 | [6] |
| **Trp•+ $^{13}C^{\varepsilon2}$** | 0.092 | [6] |
| **Trp•+ $^{13}C^{\varepsilon3}$** | 0.619 | [6] |
| **Trp•+ $^{13}C^{\zeta2}$** | -0.182 | [6] |
| **Trp•+ $^{13}C^{\zeta3}$** | -0.499 | [6] |
| **Trp•+ $^{13}C^{\eta2}$** | 0.443 | [6] |
| **Trp•+ $^{1}H^{\alpha}$** | 0 | [6] |
| **Trp•+ $^{1}H^{\beta1}$** | 2.544 | [6] |
| **Trp•+ $^{1}H^{\beta2}$** | 1.189 | [6] |
| **Trp•+ $^{1}H^{\delta1}$** | -0.421 | [6] |
| **Trp•+ $^{1}H^{\varepsilon1}$** | -0.413 | [6] |
| **Trp•+ $^{1}H^{\varepsilon3}$** | -0.504 | [6] |
| **Trp•+ $^{1}H^{\zeta2}$** | -0.050 | [6] |
| **Trp•+ $^{1}H^{\zeta3}$** | 0.124 | [6] |
| **Trp•+ $^{1}H^{\eta2}$** | -0.412 | [6] |
| **Trp•+ $^{14}N^{\varepsilon1}$** | 0.20 | [41] |
| **Trp•+ $^{15}N^{\varepsilon1}$** | 0.28 | [41] |
| **Fl•− $^{1}H^{1,8}$** | 0.3287 | [42] |
| **Fl•− $^{1}H^{2,7}$** | 0.151 | [42] |
| **Fl•− $^{1}H^{4,5}$** | 0.0889 | [42] |
| **Fl•− $^{1}H^{13}$** | 0.022 | [42] |
| **Fl•− $^{1}H^{14}$** | 0.019 | [42] |
| **Fl•− $^{1}H^{15}$** | 0.017 | [42] |
| **Fl•− $^{1}H^{16}$** | 0.009 | [42] |

[a] Scaled according to the relative hyperfine coupling constant of $^{13}C^{\alpha}$ and $^{13}C^{\gamma}$ reported in the reference and DFT-simulated hyperfine coupling constant of $^{13}C^{\gamma}$ in aqueous medium.

**Supplementary Table 3.** $^{13}C^{\alpha}$ enhancement factors (ε) and percent polarization ($P_{\%}$) of QISP Trp at 80 MHz and 600 MHz.[a]

| Concentration of QISP Trp | Enhancement factor (ε) | | Percent polarization ($P_{\%}$) | |
|---|---|---|---|---|
| | 80 MHz | 600 MHz | 80 MHz | 600 MHz |
| 5 μM | 3100.4 ± 212.2 | 432.3 ± 60.9 | 0.32 ± 0.02 | 0.33 ± 0.05 |
| 25 μM | 2183.6 ± 102.7 | 243.2 ± 28.8 | 0.24 ± 0.01 | 0.19 ± 0.02 |

[a] All data shown here were determined from experiments in Figure 2G-H and represented as average ± SE at 80 and 600 MHz (n=2, for each).

**Supplementary Table 4.** $^{1}$H enhancement factors (**ε)** of unlabeled Trp at 80 MHz and 600 MHz.

| Concentration of unlabeled Trp | Enhancement factor (ε) | | | | | |
|---|---|---|---|---|---|---|
| | 80 MHz | | | 600 MHz | | |
| | β | δ1,η3 | ε3 | β | δ1,η3 | ε3 |
| **5 μM** | -1080.8±4.0 | 194.6 ±5.2 | 473.3±1.1 | -45.6±11.3 | 35.7±6.8 | 27.4± 0.6 |
| **25 μM** | -257.0±3.0 | 66.1±1.5 | 151.3±0.7 | -46.6±1.8 | 33.9±8.0 | 25.2± 0.8 |

[a] All data shown here were derived from experiments in Fig. 3G and are shown as average ± SE at 80 and 600 MHz (n=2).

**Supplementary Table 5.** $^{1}H$ Percent polarization ($P_{\%}$) of unlabeled Trp, for data collected at 80 MHz and 600 MHz.

| Concentration of unlabeled Trp | Percent polarization ($P_{\%}$) | | | | | |
|---|---|---|---|---|---|---|
| | 80 MHz | | | 600 MHz | | |
| | β | δ1,η3 | ε3 | β | δ1,η3 | ε3 |
| **5 μM** | -0.7±0.003 | 0.1±0.004 | 0.3±0.001 | -0.3±0.008 | 0.2±0.04 | 0.1± 0.004 |
| **25 μM** | -0.2±0.002 | 0.04±0.001 | 0.1±0.001 | -0.2±0.008 | 0.2±0.003 | 0.1± 0.01 |

[a] All data shown here were derived from experiments in Fig. 3H and are shown as average ± SE at 80 and 600 MHz (n=2).

**Supplementary Table 6.** $^1$H $T_1$ relaxation times of unlabeled Trp in $D_2O$-based potassium phosphate buffer (pH ~7.2). data were collected at 80 and 600 MHz.

| Nucleus | 80 MHz [a] | 600 MHz [a] |
|---|---|---|
| $^1H^{\beta 1}$ | 0.69±0.001 | 0.71±0.025 |
| $^1H^{\beta 2}$ | 0.69±0.001 | 0.69±0.018 |
| $^1H^{\alpha 1}$ | 2.44±0.32 | 2.47±0.183 |
| $^1H^{\zeta 3}$ | 2.47±0.02 | 2.56±0.081 |
| $^1H^{\eta 2}$ | 2.97±0.11 | 2.95±0.045 |
| $^1H^{\delta 1}$ | 4.28±0.03 | 4.90±0.590 |
| $^1H^{\zeta 2}$ | 3.91±0.03 | 4.69±0.080 |
| $^1H^{\varepsilon 3}$ | 2.35±0.005 | 2.56±0.029 |

[a] The data were collected with an inversion recovery pulse sequence with no solvent suppression. All data are represented as average ± SE with n=2 at both fields.

**Supplementary Table 7.** $^1H$ $T_1$ relaxation times of unlabeled Trp in $H_2O$- and $D_2O$-based potassium phosphate buffer (pH ~7.2) at 80 MHz.

| Nucleus | In $H_2O$ [a] | In $D_2O$ [a] |
|---|---|---|
| $^1H^{\beta 1}$ | 0.52 ±0.04 | 0.69±0.001 |
| $^1H^{\beta 2}$ | 0.52 ±0.04 | 0.69±0.001 |
| $^1H^{\alpha 1}$ | 0.91±0.10 | 2.44±0.32 |
| $^1H^{\zeta 3}$ | 1.83±0.05 | 2.47±0.02 |
| $^1H^{\eta 2}$ | 1.83±0.06 | 2.97±0.11 |
| $^1H^{\delta 1}$ | 1.92±0.04 | 4.28±0.03 |
| $^1H^{\zeta 2}$ | 2.33±0.14 | 3.91±0.03 |
| $^1H^{\varepsilon 3}$ | 1.79±0.07 | 2.35±0.005 |

[a] Data were collected at 80 MHz with an inversion recovery pulse sequence in the presence (**Supplementary Fig. 10**) and absence of a WET solvent-suppression scheme, respectively. All data are presented as average ± SE, with n=4 for $H_2O$ and n=2 for $D_2O$.

#### Supplementary Figures

##### Supplementary Figure 1

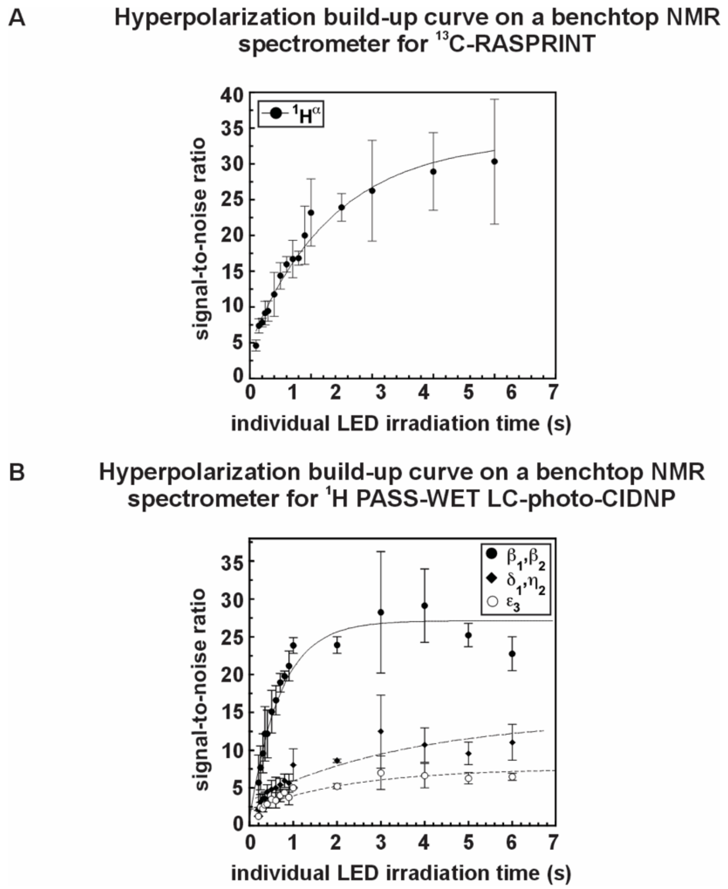


**Supplementary Figure 1. LC-photo-CIDNP experiments as a function of LED irradiation time**. **A**. Hyperpolarization build-up of the 1Ha-13Ca for 13C RASPRINT on a 100 μM QISP Trp sample (n=3). **B**. Hyperpolarization build-up as 1H PASS-WET LC-photo-CIDNP on a 100 mM unlabeled Trp sample (n=3). Three different regions of the spectrum were shown in the plot. **B.** Hyperpolarization build-up of the $^{1}H^{\alpha}$-$^{13}C^{\alpha}$ for $^{13}C$ RASPRINT on a 100 μM QISP Trp sample (n=3). A recycle delay of 1 s and 500 ms were used for $^{1}H$ PASS-WET LC-photo-CIDNP and $^{13}C$ RASPRINT experiments respectively. 25 μM fluorescein was used for all 100 μM unlabeled or QISP Trp samples. The ".sino" command in TopSpin 4.4.0 was used to measure the signal to noise

(S/N) for all the resonances shown here. All independent experiments were performed on the same day and using the same NMR tube. All data were acquired at 80 MHz (1.88 T).

#### Supplementary Figure 2

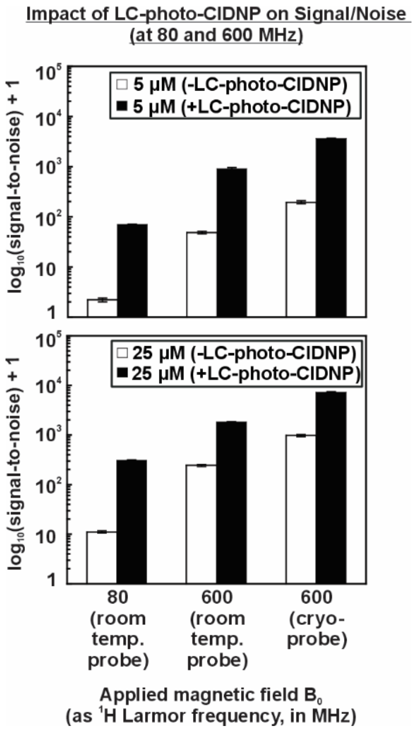


**Supplementary Figure 2. $^{13}$C RASPRINT LC-photo-CIDNP signal-to-noise (S/N) relative to non-photo-CIDNP 1D $^{1}$H NMR experiments.** The $\log_{10}$(signal-to-noise) +1 values are shown for data collected in the absence and presence of LC-photo-CIDNP at low (80 MHz, 1.88 T) and high (600 MHz, 14.1 T) field at 5 µM (top panel) and 25 µM (bottom panel) sample concentrations. For the non–photo-CIDNP condition, the S/N was first measured using a standard 1D $^{1}$H experiment on a 0.1% ethylbenzene sample in $CDCl_3$ with 32 scans, a 2 s recycle delay at both fields. The S/N obtained was then linearly scaled by concentration to estimate the expected S/N at 25 µM and 5 µM (n = 2, at each $B_0$). In the case of LC-photo-CIDNP data, the S/N values were obtained from $^{13}$C RASPRINT experiments performed under light (LED-on) conditions with 32 scans, a 0.5 s recycle delay, and 1 s LED irradiation/scan at both fields (n = 2, for each concentration and each $B_0$). QISP Trp samples at 25 µM and 5 µM were prepared with 12 µM and

8 μM fluorescein, respectively, in 10 mM potassium phosphate buffer (pH ~7.2). Predicted S/N values for the 600 MHz system lacking a cryoprobe were estimated by dividing the experimentally measured cryoprobe S/N by a factor of four, consistent with the literature.[43] A constant offset of 1 was added to the Y axis values, to avoid negative values on the $\log_{10}$ scale.

Supplementary Figure 3

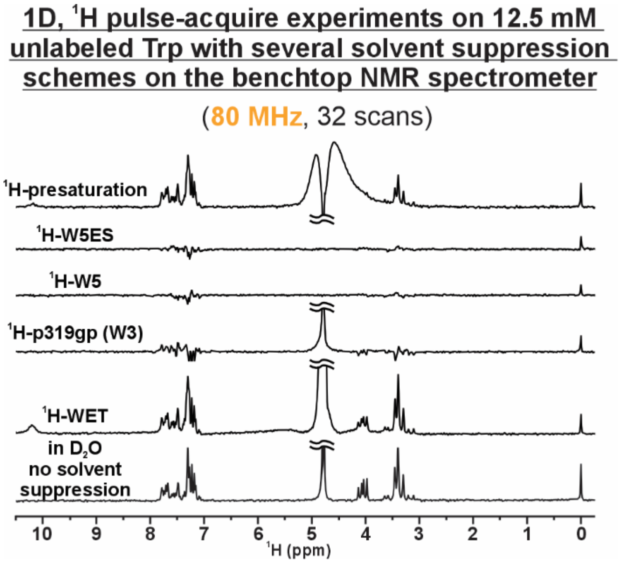


**Supplementary Figure 3. $^{1}$H-WET solvent suppression scheme produces the best result at low mM sample concentration, on a benchtop NMR spectrometer.** $^{1}$H spectra were acquired on a 12.5 mM unlabeled Trp sample in $H_2O$ using various solvent suppression techniques and compared to a reference spectrum collected in $D_2O$ (no suppression applied). Among the tested schemes, $^{1}$H-WET provided the most effective solvent suppression with a narrow suppression bandwidth, preserving the aliphatic signals. Notably, the exchangeable proton on the indole pyrrole ring remained clearly visible with $^{1}$H-WET, unlike in other methods. Based on these experiments, $^{1}$H-WET was particularly selected for all $^{1}$H-photo-CIDNP experiments on the Benchtop NMR. All samples were prepared with 500 µM DSS as the internal standard (0 ppm). All spectra were recorded with a recycle delay of 1 s.

#### Supplementary Figure 4

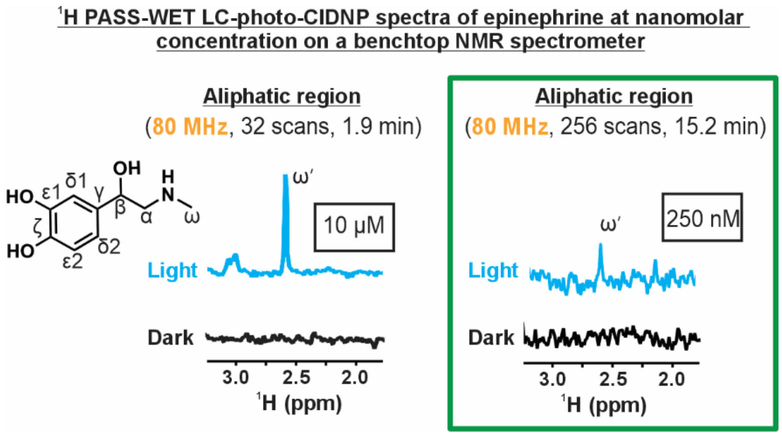


**Supplementary Figure 4. Detection of epinephrine at nM concentrations on a benchtop NMR spectrometer.** The spectra display the aliphatic region of $^1$H PASS-WET LC-photo-CIDNP data collected on 10 µM and 250 nM epinephrine in aqueous buffer. The resonance labeled (ω′) likely originates from a photoproduct of epinephrine that is strongly enhanced by LC-photo-CIDNP.[24] This photoproduct was not characterized in the present study. An LED irradiation time of 1 s per scan and a recycle delay of 1 s were used for all experiments. All data were collected at 80 MHz (1.88 T).

#### Supplementary Figure 5

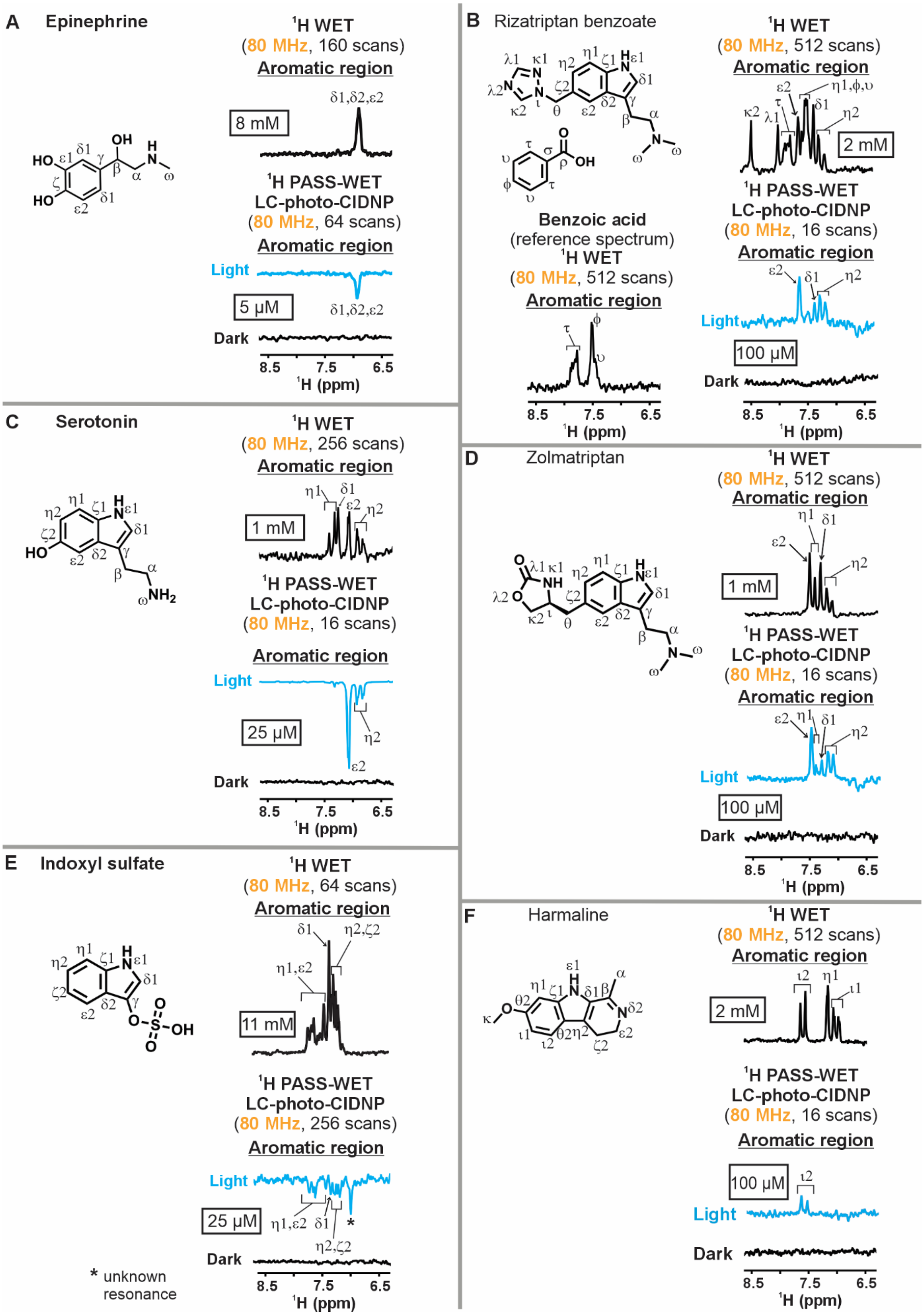

**Supplementary Figure 5. Comparison between conventional NMR (non-photo-CIDNP) and $^1$H PASS-WET LC-photo-CIDNP NMR spectra acquired on a benchtop NMR spectrometer (Part – I).** A. Epinephrine hydrochloride (racemic mixture), B. Rizatriptan benzoate (top right) and benzoic acid (bottom left), C. Serotonin, D. Zolmitriptan, E. Indoxyl sulfate, F. Harmaline. All experiments used a 1 s recycle delay and 1 s LED irradiation time per scan. Note, for harmaline, the LC-photo-CIDNP sample used here also included 40% DMSO-$d_6$ for an improved S/N, especially in the aromatic region **(Supplementary Fig. 7)**. In addition, the emissive aliphatic resonances ($^1H^{\alpha}$) for harmaline were also visible under light conditions at 100 μM but the chemical shifts were gradually going upfield with increasing DMSO-$d_6$ percentage in the sample (data not shown). All LC-photo-CIDNP samples with 25 μM and 100 μM indole-containing compounds included 12 μM and 25 μM fluorescein, respectively. LC-photo-CIDNP sample of 5 μM epinephrine included 5 μM ATTO Thio 12. All 1D, $^1$H WET spectra (no-photo-CIDNP) experiments were carried out in 10 mM potassium phosphate buffer at pH ~7.2. All data were collected at 80 MHz (1.88 T) with n=2.

#### Supplementary Figure 6

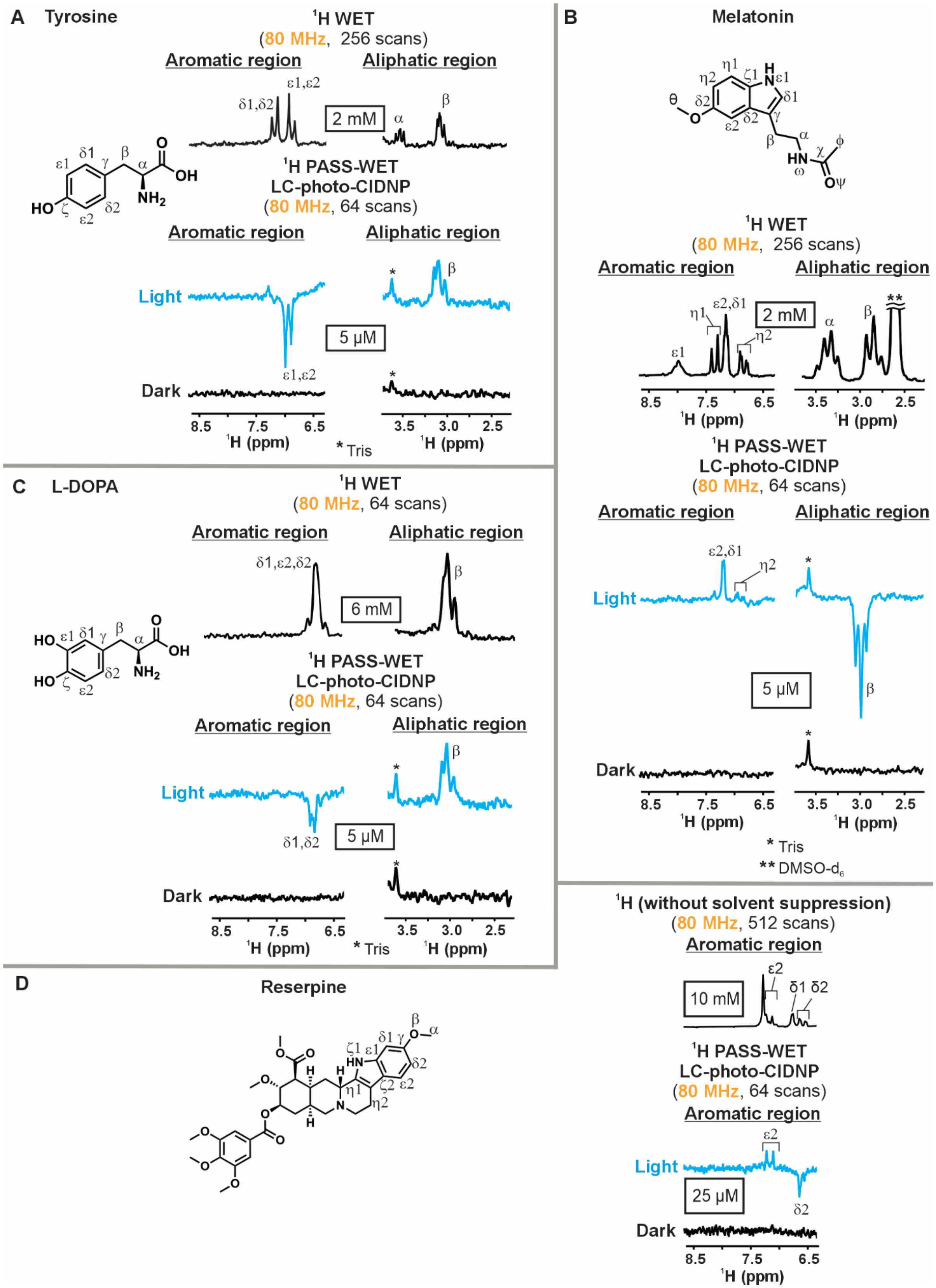

**Supplementary Figure 6. Comparison between conventional NMR (non-photo-CIDNP) and $^{1}$H PASS-WET LC-photo-CIDNP NMR spectra acquired on a benchtop NMR spectrometer (Part – II).** A. Tyrosine, B. Melatonin, C. L-DOPA, D. Reserpine. All experiments used a 1 s recycle delay and 1 s LED irradiation time per scan. All LC-photo-CIDNP samples with 25 μM and 5 μM indole-containing compounds included 12 μM and 8 μM fluorescein, respectively. All LC-photo-CIDNP samples with 5 μM phenolic compounds included 5 μM ATTO Thio 12. All 1D, $^{1}$H WET spectra (no-photo-CIDNP) experiments were carried out in 10 mM potassium phosphate buffer at pH ~7.2, except reserpine. For a 10 mM reserpine sample, a 1D, $^{1}$H experiment was carried out in DMSO-$d_6$ owing to its extremely poor solubility in water-based buffer. In addition, a ~0.04 ppm upfield shift was observed in the LC-photo-CIDNP spectrum of 25 μM reserpine under light conditions compared to the 10 mM 1D, $^{1}$H spectrum. All data were collected at 80 MHz (1.88 T) with n=2.

### Supplementary Figure 7

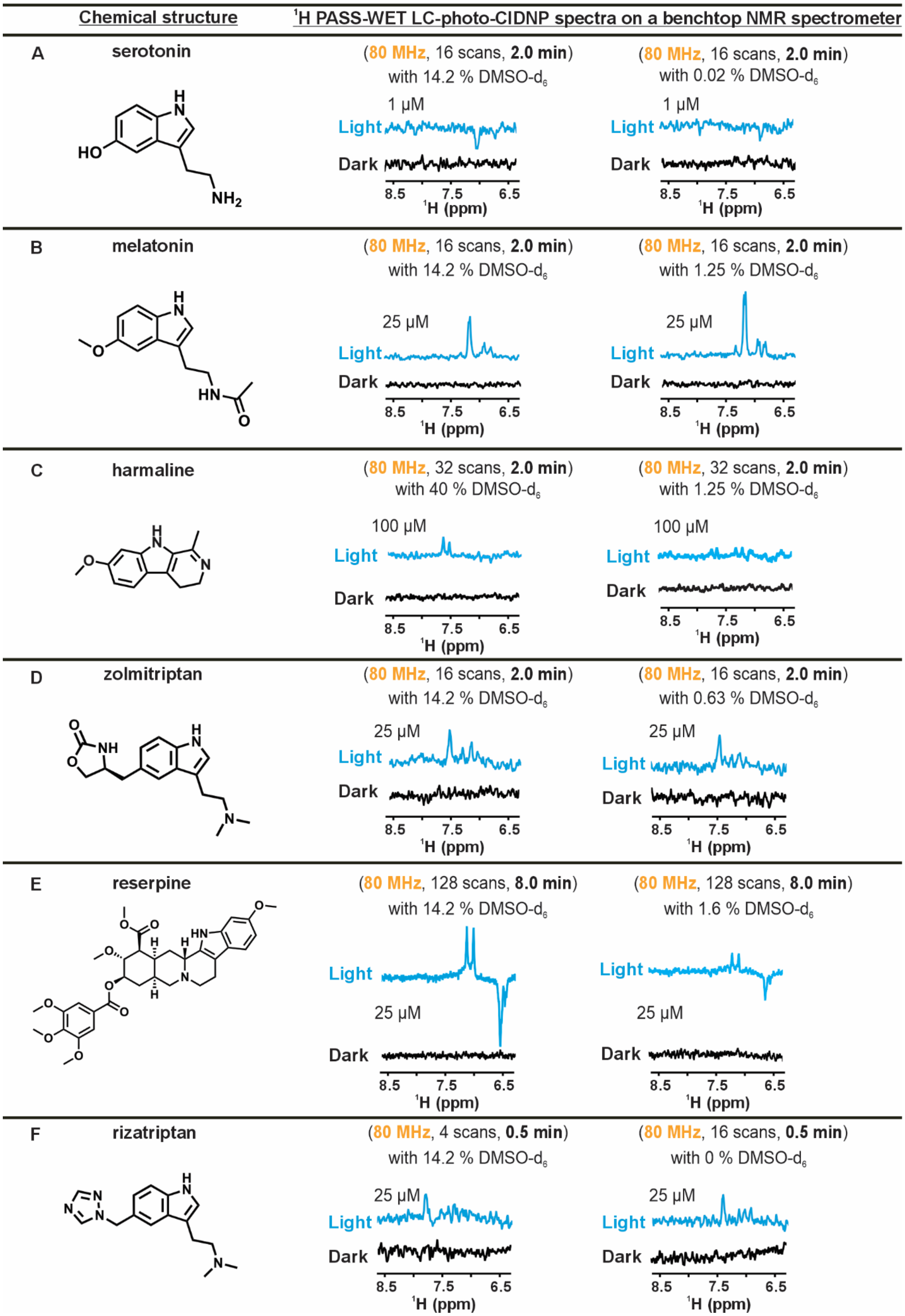

**Supplementary Figure 7. $^1H$ PASS-WET LC-photo-CIDNP data collected in the absence and presence of DMSO on neurotransmitters and pharmaceuticals.** $^1H$ PASS-WET LC-photo-CIDNP spectra collected on A. serotonin (16 scans), B. Melatonin (16 scans), C. Harmaline (32 scans), D. Zolmitriptan (16 scans), E. Reserpine (128 scans) and, F. Rizatriptan benzoate (16 scans) with additional DMSO-$d_6$ (left column) and without additional DMSO-$d_6$ (right column). We noted only for harmaline and reserpine especially, addition of DMSO-$d_6$ improved the S/N and for the rest, there was not a significant difference. A 1 s recycle delay and 1 s LED irradiation per scan were used for all experiments. All data were acquired at 80 MHz (1.88 T) with n=2.

#### Supplementary Figure 8

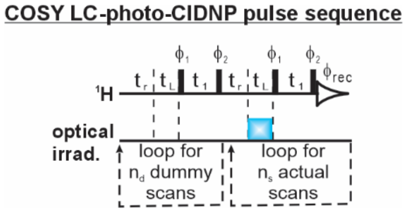


**Supplementary Figure 8. COSY LC-photo-CIDNP pulse sequence used in this study.** The first and second halves correspond to the dummy ($n_d$) and actual scans ($n_s$), respectively. The two halves are identical except that LED irradiation is applied only during the actual scan. $t_r$ and $t_L$ represent recycle delay and LED irradiation time, respectively. During our experiments, $n_d$ was set to 16. The phases used for this sequence are the following: $\phi_1$ =x, x, x, x, y, y, y, y, y, -x, -x, -x, -x, -y, -y, -y, -y; $\phi_2$ = x, y, -x, -y; $\phi_{rec}$ = x, -x, x, -x, -y, y, -y, y, -x, x, -x, x, y, -y, y, -y.

#### Supplementary Figure 9

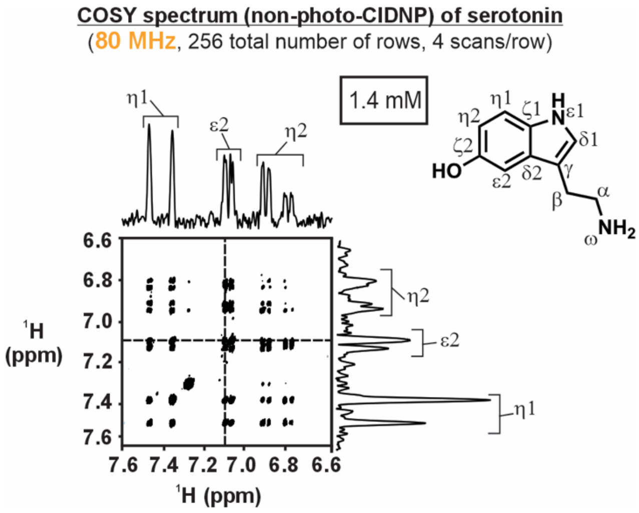


**Supplementary Figure 9. Conventional (non-photo-CIDNP) COSY NMR spectrum of serotonin.** COSY NMR of 1.4 mM serotonin in aqueous buffer was collected with 256 total rows and 4 scans/row. The dashed lines indicate the position of the 1D $^{1}$H spectral slices displayed on top of the 2D spectrum. See methods for experimental details. This data was collected at 80 MHz (1.88 T).

#### Supplementary Figure 10

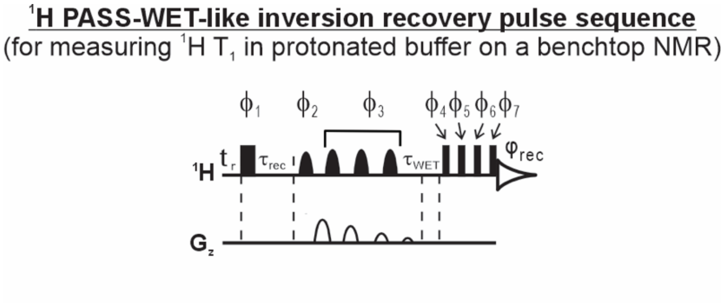


**Supplementary Figure 10. $^1$H PASS-WET-like inversion recovery sequence used in this study for measuring $^1$H $T_1$ in protonated buffer.** $\tau_r$, $\tau_{rec}$ and $\tau_{WET}$ represent recycle delay, inversion recovery delay and WET delay, respectively. The phase cycling is $\phi_{rec}$ is x, -x, x, -x, y, -y, y, -y; $\phi_1 =$ x, -x; $\phi_2 =$ x; $\phi_3$ = y, $\phi_4$ = x, y, y, x, -x, x, x, -x; $\phi_5$ =-x, x, -x, x, -y, y, -y, y; $\phi_6 =$ -y, y, y, -y, x, -x, -x, x; $\phi_7 =$ x, -x, x, -x, y, -y, y, -y. This pulse sequence was only used at 80 MHz (1.88 T).

**Supplementary Figure 11**

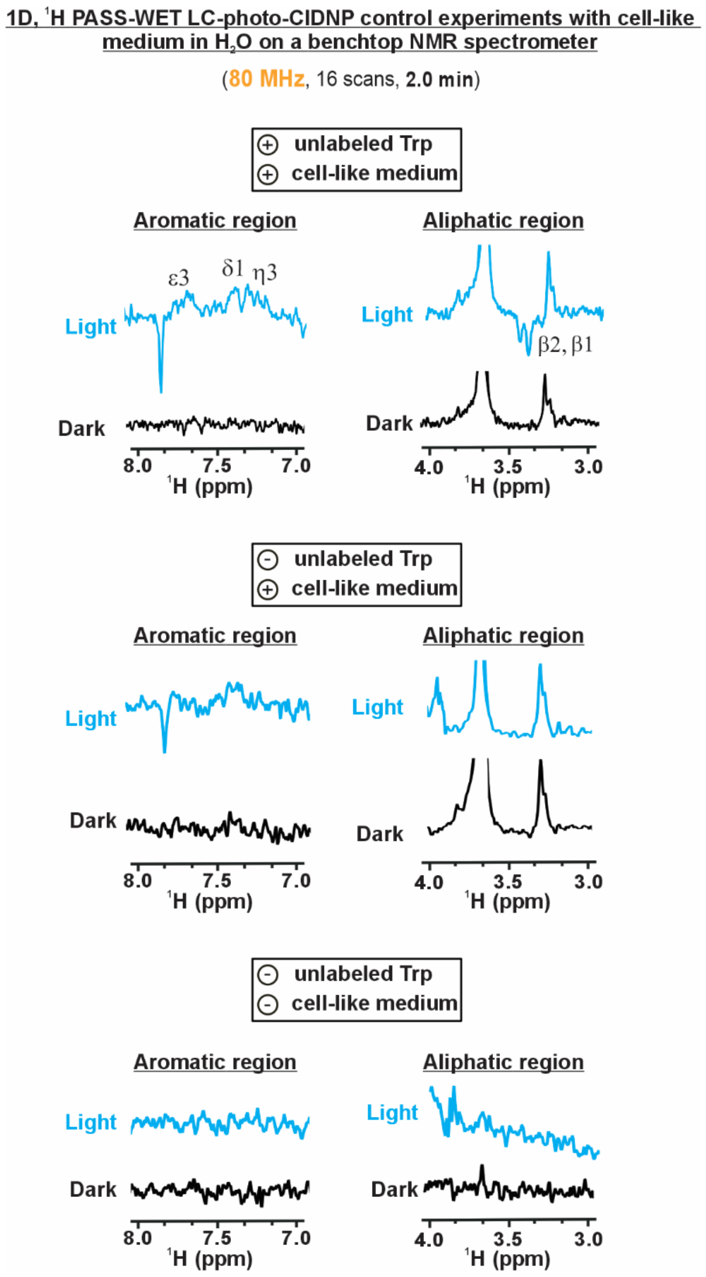


**Supplementary Figure 11**. **Control experiments using ¹H PASS-WET LC-photo-CIDNP in *E. coli* cell-like medium.** ¹H PASS-WET LC-photo-CIDNP spectra collected on a sample containing *E. coli* cell-like medium in 10 mM aqueous buffer with all LC-photo-CIDNP components, with 25 μM tryptophan, under light and dark conditions (top). ¹H PASS-WET LC-photo-CIDNP spectra collected on a sample containing all LC-photo-CIDNP components, with *E.*

*coli* cell-like medium but without Trp at natural abundance, under light and dark conditions (middle). $^1$H PASS-WET LC-photo-CIDNP spectra collected on a sample containing all components without unlabeled Trp and without *E. coli* cell-like medium (bottom). The absence of strong emissive resonances around 3.5 ppm and weak absorptive resonances in the aromatic region in samples lacking unlabeled Trp confirmed that these signals originated from the $^1H^\beta$ protons and aromatic protons of Trp, respectively. In contrast, the persistent emissive resonance at ~7.8 ppm, even without Trp, suggested it came from endogenous *E. coli* components enhanced by LC-photo-CIDNP. All $^1$H PASS-WET LC-photo-CIDNP spectra were collected with a recycle delay of 1 s and an LED irradiation time of 6 s per scan and used 25 μM fluorescein. All data were acquired at 80 MHz (1.88 T).

#### Supplementary Figure 12

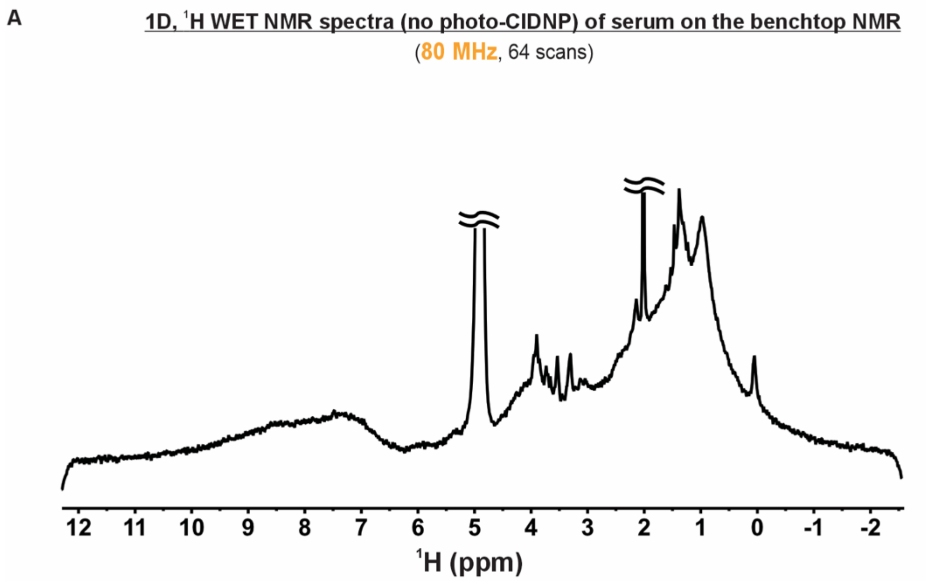


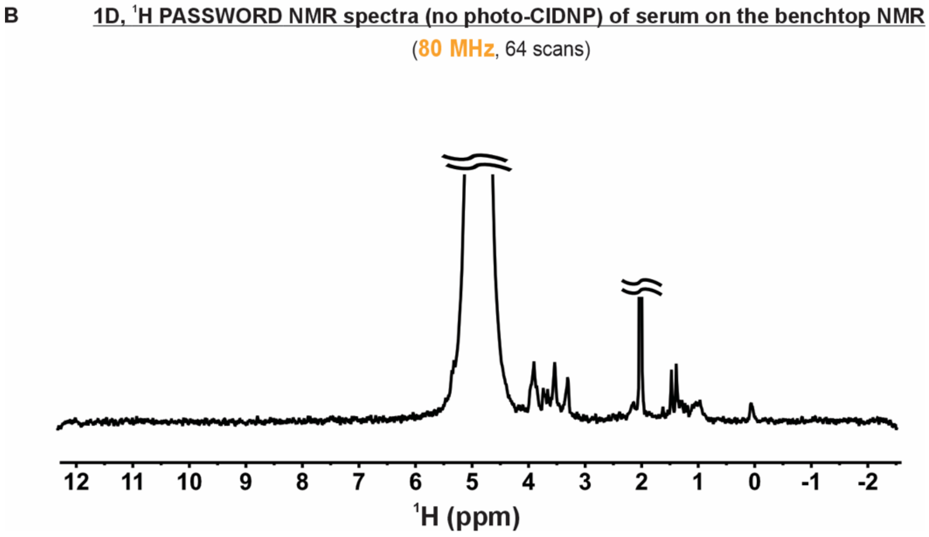


**Supplementary Figure 12. $^1$H PASSWORD in serum offers better background suppression than $^1$H WET experiment. A.** $^1$H WET and **B.** $^1$H PASSWORD experiment on a 2-times diluted serum sample. $^1$H WET-CPMG uses n = 126 $T_2$ filtering loops (corresponding to 79 ms). Both the experiments used 64 scans and 1 s of recycle delay. All data were acquired at 80 MHz (1.88 T).

#### Supplementary Figure 13

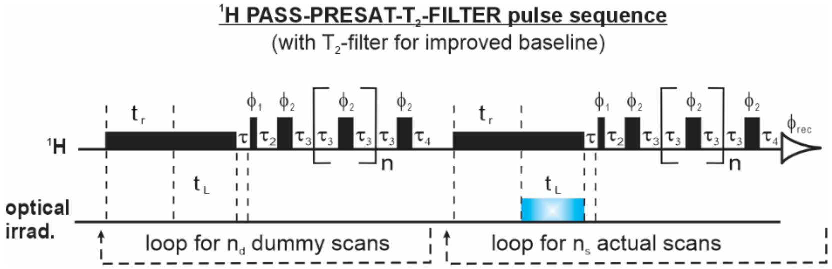


**Supplementary Figure 13. $^1$H PASS-PRESAT-T$_2$-FILTER pulse sequence used in this work**. The first and second halves correspond to the dummy and acquisition scans, respectively. The two halves are identical except that LED irradiation is applied only during the actual scan. $t_r$ and $t_L$ represent recycle delay and LED irradiation time, respectively. A 3 s presaturation delay was used with a power of 50 Hz. LED was gated on during the last 1 s of the presaturation pulse to minimize the water-signal recovery during LED irradiation. For these experiments, $n_d$ was set to 4. The T$_2$-filter loop count (n) was set to 126. The phase cycle used for this sequence: $\phi_1$ = x, x, -x, -x, y, y, -y, -y; $\phi_2$ = y, -y, y, -y, x, -x, x, -x; $\phi_{rec}$ = x, x, -x, -x, y, y, -y, -y. This pulse sequence was only used at 600 MHz (14.1 T).

#### Supplementary Figure 14

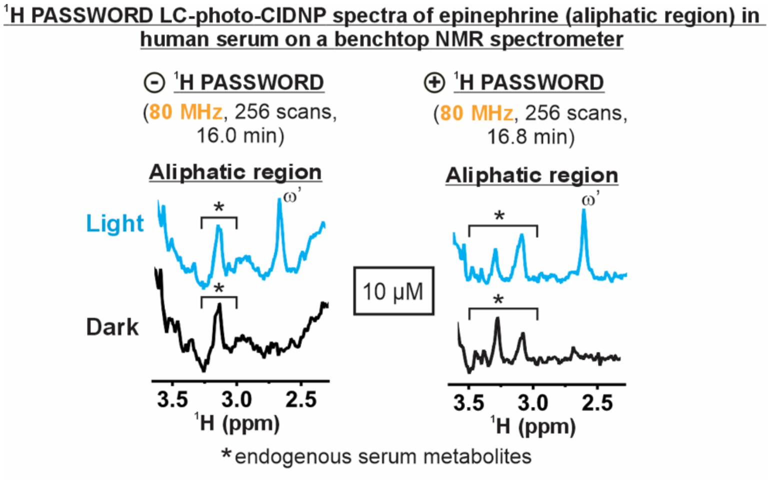


**Supplementary Figure 14. LC-photo-CIDNP enables low-micromolar detection of neurotransmitters (epinephrine) in human serum on the benchtop NMR.** LC-photo-CIDNP spectra of 10 μM epinephrine in serum under light and dark conditions without and with $^{1}$H PASSWORD experiment. The photoproduct of epinephrine (ω') was revealed under light (LED-on) conditions. An LED irradiation time of 1 s per scan and a recycle delay of 1 s were used for all experiments. All spectra were acquired at 80 MHz (1.88 T).

#### Supplementary Figure 15

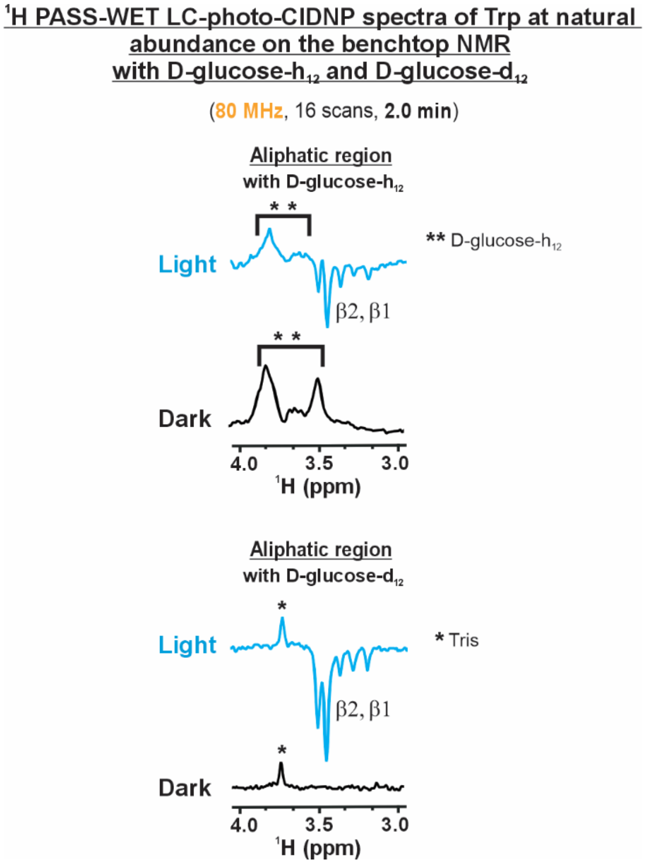


**Supplementary Figure 15. $^1$H PASS-WET LC-photo-CIDNP with D-glucose-d$_{12}$ improved the S/N in the aliphatic region.** Top: Spectra acquired for 25 µM unlabeled Trp with D-glucose-h$_{12}$ under light and dark conditions. Bottom: Same experiment performed with D-glucose-d$_{12}$. With glucose-d$_{12}$, the entire $^1$H$^\beta$ region is clearly visible, free from interfering signals. In contrast, with glucose-h$_{12}$, the $^1$H$^\beta$ region is partially obscured due to the intense background resonances from D-glucose-h$_{12}$ and D-glucono-1,5-lactone - latter one formed as a byproduct of the reaction between glucose-h$_{12}$, glucose oxidase, and catalase. The use of D-glucose-d$_{12}$ leads to the formation of a deuterated D-glucono-1,5-lactone species, which produces almost negligible signal in the aliphatic region (3.2–4.2 ppm). A resonance at ~3.7 ppm was noticeable even with D-glucose-d$_{12}$; which originated from Tris buffer present in the glucose oxidase powder, not from residual non-deuterated glucose species. This was confirmed by control experiments with only glucose oxidase (data not shown). A 1 s recycle delay, and 6 s LED irradiation per scan were used for $^{1}H$ PASS-WET LC-photo-CIDNP. Both experiments contained 12 μM fluorescein. All data were acquired at 80 MHz (1.88 T).

#### Supplementary Figure 16

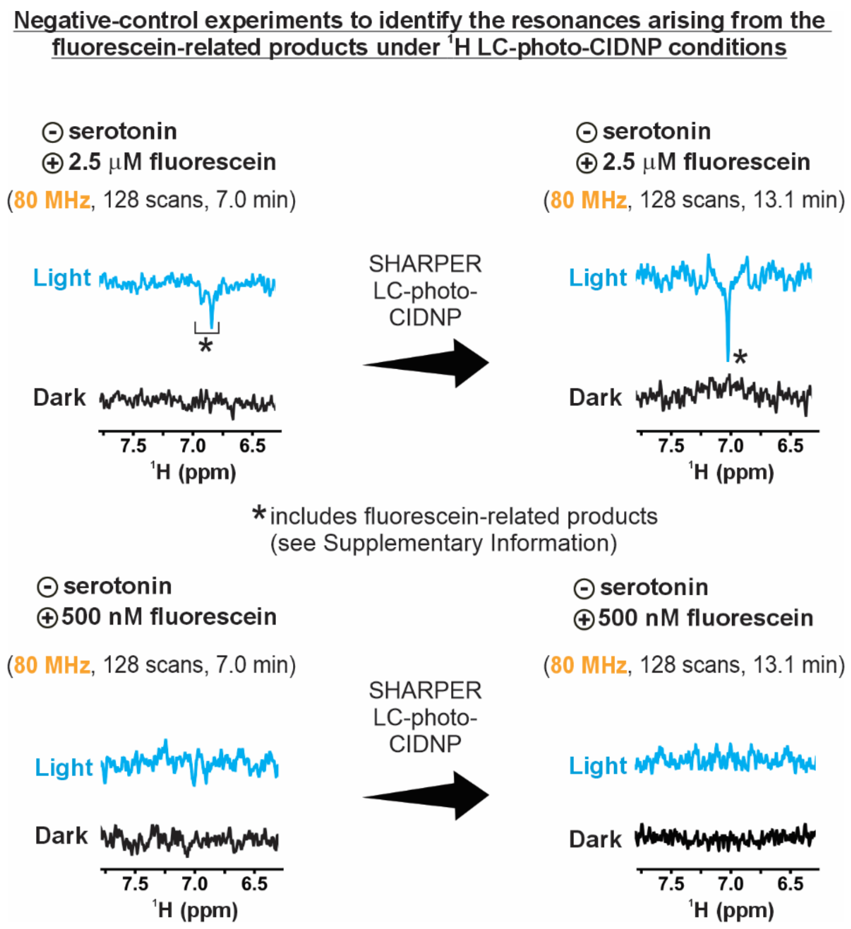


**Supplementary Figure 16. Control ¹H LC-photo-CIDNP experiments for identifying fluorescein-derived resonances.** Top: ¹H LC-photo-CIDNP spectra acquired without SHARPER (left) and with SHARPER (right) on a sample containing 2.5 μM fluorescein and all LC-photo-CIDNP components except the molecule of interest. Bottom**:** ¹H LC-photo-CIDNP spectra acquired without SHARPER (left) and with SHARPER (right) on a sample containing 500 nM fluorescein and all LC-photo-CIDNP components except the molecule of interest. A small number of resonances arising from fluorescein-related products were observed. However, characterization of these species is beyond the scope of this work. An LED irradiation time of 1 s per scan and a recycle delay of 1 s were used for all experiments. All spectra were acquired at 80 MHz (1.88 T).

### Appendix

**Python script used to generate NMR spectra devoid of the contributions due to the presence of a cryogenic probe**

```
import numpy as np
import matplotlib.pyplot as plt
import os

file_path = input("Full path to my spectrum file (.txt): ").strip()

spectra_name_base = input(
    "Enter a name for the output files (e.g., '5uM_QISP_32_scans'): "
).strip()

ppm, intensity = np.loadtxt(file_path, unpack=True)

# Typical cryoprobe sensitivity enhancement
cryoprobe_enhancement_factor = 3.5


# Use the baseline between 8 and 10 ppm to estimate the noise
baseline_indices = np.where((ppm >= 8.0) & (ppm <= 10.0))[0]
noise_std = np.std(intensity[baseline_indices])


# Add noise to simulate spectra with and without a cryoprobe
noise_with_cryo = np.random.normal(
    0, 6 * noise_std, size=len(intensity)
)

noise_without_cryo = np.random.normal(
    0,
    6 * noise_std * cryoprobe_enhancement_factor,
    size=len(intensity)
)

noisy_intensity_with_cryo = intensity + noise_with_cryo
noisy_intensity_without_cryo = intensity + noise_without_cryo


# Plot the simulated noise and spectra
plt.figure(figsize=(12, 6))
```

```
plt.subplot(1, 2, 1)

plt.plot(
    ppm,
    noise_with_cryo,
    label="Noise with Cryoprobe",
    color="blue",
    alpha=0.7
)

plt.plot(
    ppm,
    noise_without_cryo,
    label="Noise without Cryoprobe",
    color="red",
    alpha=0.7
)

plt.gca().invert_xaxis()
plt.xlabel("Chemical Shift (ppm)")
plt.ylabel("Noise Intensity")
plt.title("Noise Comparison")
plt.legend()


plt.subplot(1, 2, 2)

plt.plot(
    ppm,
    noisy_intensity_with_cryo,
    label="With Cryoprobe",
    color="blue"
)

plt.plot(
    ppm,
    noisy_intensity_without_cryo,
    label="Without Cryoprobe",
    color="red"
)

plt.gca().invert_xaxis()
plt.xlabel("Chemical Shift (ppm)")
plt.ylabel("Intensity")
plt.title("Spectra Comparison")
plt.legend()
```

```python
plt.tight_layout()
plt.show()


# Output file names
output_file_with_cryo = (
    f"{spectra_name_base}_with_cryo_simulated_light.txt"
)

output_file_without_cryo = (
    f"{spectra_name_base}_without_cryo_simulated_light.txt"
)

output_noise_with_cryo = f"{spectra_name_base}_noise_with_cryo.txt"
output_noise_without_cryo = f"{spectra_name_base}_noise_without_cryo.txt"


# Save the spectra
np.savetxt(
    output_file_with_cryo,
    np.column_stack((ppm, noisy_intensity_with_cryo))
)

np.savetxt(
    output_file_without_cryo,
    np.column_stack((ppm, noisy_intensity_without_cryo))
)


# Save the noise separately
np.savetxt(
    output_noise_with_cryo,
    np.column_stack((ppm, noise_with_cryo))
)

np.savetxt(
    output_noise_without_cryo,
    np.column_stack((ppm, noise_without_cryo))
)


print("Files saved:")
print(output_file_with_cryo)
print(output_file_without_cryo)
```

```
print(output_noise_with_cryo)
print(output_noise_without_cryo)
```
